# A new species of *Cichlidogyrus* Paperna, 1960 (Platyhelminthes: Monogenea: Dactylogyridae) infecting tilapias in Lake Kariba (Zimbabwe), with a discussion on its phylogenetic position

**DOI:** 10.1101/2022.05.29.493922

**Authors:** Mare Geraerts, Tine Huyse, Maarten P. M. Vanhove, Tom Artois

## Abstract

Monogeneans dominate the external parasite fauna of bony fish. During recent years, examination of more populations of species of *Cichlidogyrus* Paperna, 1960 has led to the (re)description of several species. *Cichlidogyrus halli* (Price & Kirk, 1967) Price, 1968, for example, has been redescribed several times in the past and has been proposed to encompass many (pseudo)cryptic species. In Lake Kariba (Zimbabwe), specimens of a species of *Cichlidogyrus* were found that morphologically resemble *C. halli*. These specimens were found on the gills of native *Oreochromis* cf. *mortimeri* and *Coptodon rendalli* (Boulenger, 1897), and introduced Nile tilapia *Oreochromis niloticus* (Linnaeus, 1758). A detailed study of the morphology of these specimens, including morphometrics, and a thorough comparison with specimens of *C. halli* is presented. Part of the *COI* gene and 18S-ITS1 fragment were sequenced and analysed to provide insight into the phylogenetic placement of these specimens within the *Cichlidogyrus-Scutogyrus* monophylum. We found that *C. halli* and the new specimens sp. nov. are sister clades within the same monophyletic clade, and that clear morphological and morphometric differences are present in the dorsal bar of the haptor and accessory piece of the male copulatory organ. Based on these results, the new specimens are described as a new species: *C. chloeae* sp. nov. The role of introduced Nile tilapia as a potential reservoir for native parasites raises concern for potential spillbacks and stresses the need for further monitoring of monogeneans on native and introduced tilapias.

## Introduction

Monogenea is a taxon of Platyhelminthes, dominating the external parasite fauna of bony fish (Cribb et al., 2002; Paladini et al., 2017; Pugachev et al., 2010). It is a group of small hermaphrodite flatworms (ranging from ca. 100 µm to 4 cm) with a direct life cycle and with most species being host specific (Paladini et al., 2017; Řehulková et al., 2018). Species identification is traditionally based on the morphology of the sclerotised parts of the posterior attachment organ, called (opist)haptor, and the male copulatory organ (MCO) and vagina. (e.g. Pariselle & Euzet 2009). Among African monogeneans, *Cichlidogyrus* Paperna, 1960 is the most speciose genus (Pariselle & Euzet, 2009; Řehulková et al., 2018) with currently 128 described species from a total of 117 African cichlid species (Cruz-Laufer et al., 2021a).

Over the years, the examination of more populations of *Cichlidogyrus* spp. has led to the recent (re)description of several species (Fannes et al., 2017; Gobbin et al., 2021; Igeh et al., 2017; Jorissen et al., 2018b; Pariselle et al., 2003). Additionally, several species of Monogenea, including species of *Cichlidogyrus*, have been reported to display intraspecific morphological variability correlated with host species and geographic distribution (Kmentová et al., 2018; Rahmouni et al., 2021). *Cichlidogyrus halli* (Price & Kirk, 1967) Price, 1968, for example, is a morphologically variable species having been redescribed several times in the past (El-Naggar & Khidr, 1985; Ergens, 1981). Moreover, several morphotypes (e.g. Jorissen et al. 2018a) and subspecies (e.g. Paperna, 1979) within this species have been proposed, though the conspecific status of these subspecies has been questioned (Douëllou, 1993; Jorissen et al., 2018a; Jorissen et al., 2021; Pouyaud et al., 2006).

In the 1990s, Douëllou (1993) examined specimens of *C. halli* infecting cichlids in Lake Kariba (Zimbabwe) and found that their morphology deviates from the one in the original species description in having longer auricles. However, she refrained from describing them as a separate taxon (Douëllou, 1993). During a field expedition in 2019, specimens of ‘*C. halli*’, morphologically similar to the specimens reported by Douëllou (1993), were found infecting several tilapia species present in Lake Kariba.^1^

Lake Kariba is a man-made lake created in 1958 by damming the fast flowing middle Zambezi River (Reeve, 1960). Only three tilapia species are indigenous in the middle Zambezi Basin: *Oreochromis mortimeri* (Trewavas, 1966), *Coptodon rendalli* (Boulenger, 1897), and *Tilapia sparrmanii* Smith, 1840 (Marshall, 1988; Skelton, 1993). Nile tilapia, *Oreochromis niloticus* (Linnaeus, 1758), has been introduced for aquaculture purposes and has become the most dominant tilapia species in the lake (Froese & Pauly, 2021; Maulu & Musuka, 2018).

In the present study, the re-evaluation of additional specimens of ‘*C. halli*’, and the morphological and genetical comparison of these specimens with *C. halli*, have led to the description of a new species of *Cichlidogyrus*, namely *C. chloeae* sp. nov.

## Material and methods

### Collection, sample preparation and conservation

During a field expedition at Lake Kariba in October–November 2019, tilapias were purchased from local fishermen, who caught the fish in the lake by drift netting. These specimens belong to three different species: *O. niloticus, C. rendalli*, and *O*. cf. *mortimeri*. For details about the sampling locations, we refer to **Fig. 1** and **Table 1**. The identification of the specimens resembling *O. mortimeri* (Trewavas, 1966) is uncertain as they show the enlarged jaws typical of *O. mossambicus*, which could point towards hybridisation. Additionally, several specimens of *O. niloticus* were bought at two local fish farms, Lake Harvest and Nicholson Bream Farm, located nearby the lake (**Fig. 1**; **Table 1**). These fish were caught by seine netting. In case they were still alive, fish were killed by severing the spinal cord. Fish were morphologically identified in the field. From each specimen, a fin clip was taken and stored in 99% (v/v) ethanol for later genetic identification. The gills from both gill chambers were dissected and stored in 99% (v/v) ethanol. In the laboratory, the gills were exhaustively screened for monogeneans using a Nikon C-DS stereomicroscope and an entomological needle. Some monogeneans were mounted for morphological examination on a glass slide, fixed with lactophenol, and covered with a coverslip. Coverslips were sealed with kolophonium-lanoline wax. The remaining flatworms were stored in 99% (v/v) ethanol for genetic identification.

**Fig. 1.**
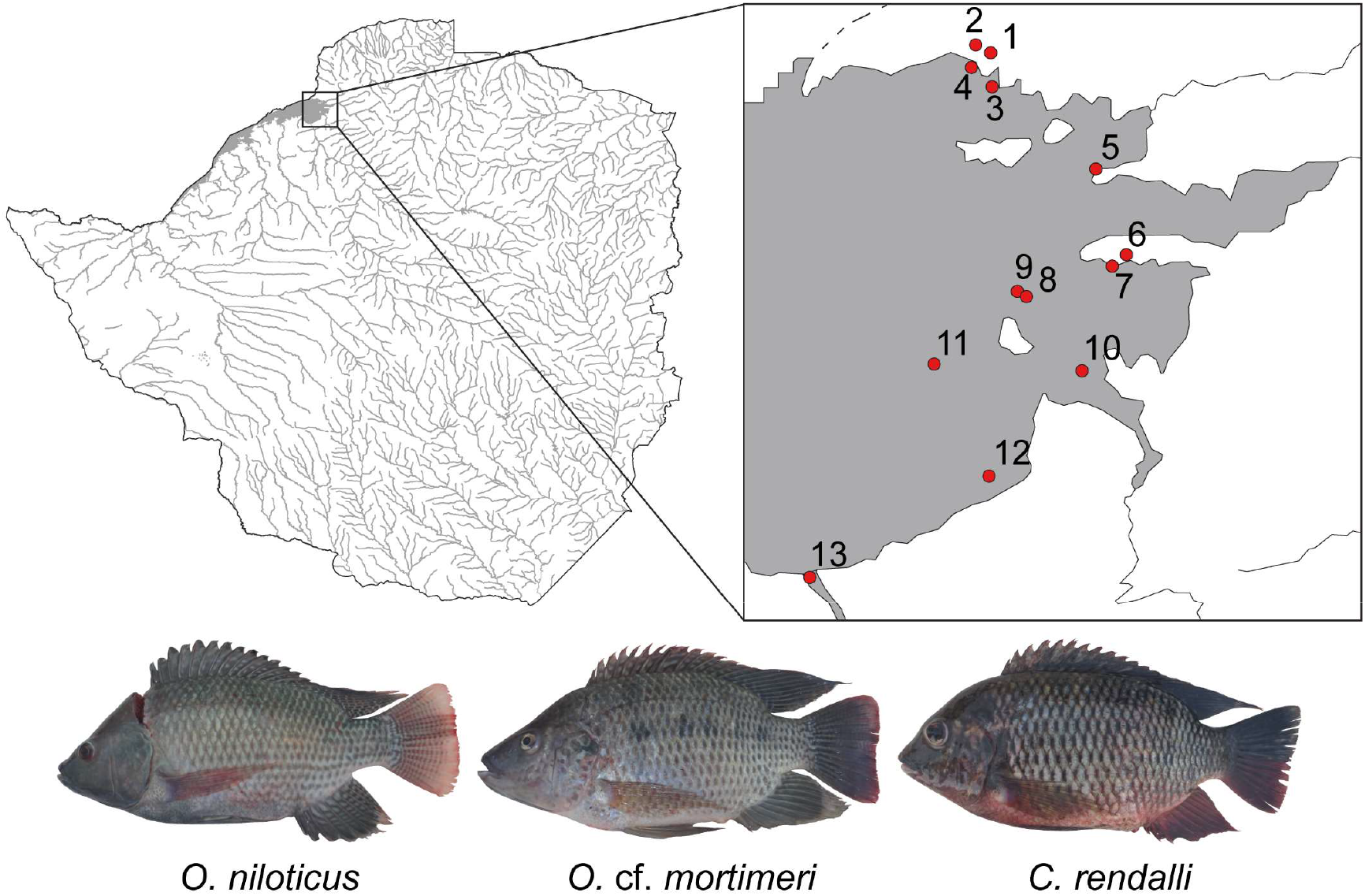
Map of Zimbabwe on the left with the framed region expanded on the right. Red dots indicate different sampling localities. The numbers of the localities correspond with those in **Table 1**. At the bottom the three tilapia species that were collected in the present study.

**Table 1.** Sampling locations in Lake Kariba and in farms near Lake Kariba with the location label, details of the sampling location, and coordinates. Location labels correspond to the ones in **Fig. 1**.

| Location label | Details | Longitude | Latitude |
| --- | --- | --- | --- |
| 1 | Nicholson Bream Farm | −16.5304 | 28.8608 |
| 2 | Lake Harvest Aquaculture | −16.5259 | 28.8524 |
| 3 | Lake Kariba at discharge channel of crocodile farm | −16.5495 | 28.8616 |
| 4 | Green Water in Lake Kariba | −16.5386 | 28.8500 |
| 5 | Fishermen village Gache Gache Cooperation at Lake Kariba | −16.5956 | 28.9198 |
| 6 | Gatche River Bay in Lake Kariba | −16.6437 | 28.9369 |
| 7 | Gatche River Bay in Lake Kariba | −16.6502 | 28.9290 |
| 8 | Gatche River Bay in Lake Kariba | −16.6672 | 28.8809 |
| 9 | Open water in Lake Kariba | −16.6643 | 28.8758 |
| 10 | Gatche River Bay in Lake Kariba | −16.7088 | 28.9121 |
| 11 | Open water in Lake Kariba | −16.7051 | 28.8291 |
| 12 | Fishermen at Gache Gache | −16.7680 | 28.8599 |
| 13 | Sayati Gorge | −16.8248 | 28.7592 |

Fin clips were deposited in the ichthyology collection at the Royal Museum for Central Africa (RMCA) in Tervuren (Belgium) under the collection number RMCA 2022.007.P. Mounted parasite specimens were deposited in the invertebrate collection of the RMCA; the collection of the research group Zoology: Biodiversity and Toxicology at Hasselt University, Diepenbeek, Belgium (HU); and the Finnish Museum of Natural History, Helsinki, Finland (MZH) (see ‘HOLOTYPE’ and ‘PARATYPE’ in the Results section for details on repositories and accession numbers).

### Microscopy and illustrations

The whole-mounts were examined under a Leica DM2500 microscope using differential interference contrast (DIC). Species were identified to genus level following the identification key in Řehulková et al. (2018) and to species level with the identification key in Pariselle & Euzet (2009). Throughout this paper, we follow the terminology, the method of measuring the different parts of the sclerites, and the numbering of the uncinuli as in Geraerts et al. (2020). Species descriptions are focused on details of the sclerotised parts i.e. haptor, male copulatory organ (MCO) and vagina (if sclerotised). Additional measurements were taken for the ventral and dorsal bar to enable a morphological comparison with previous studies (**Fig. 2**). Measurements and photographs were taken with the Leica Application Suite X (LASX) software. Drawings were made freehand using a drawing tube at a magnification of 1000× (objective ×100 immersion, ocular ×10) and edited in Adobe Illustrator version 25.2.3. Drawings of the different sclerotised parts were based on multiple specimens in case not all structures were clearly visible in a single individual.

**Fig. 2.**
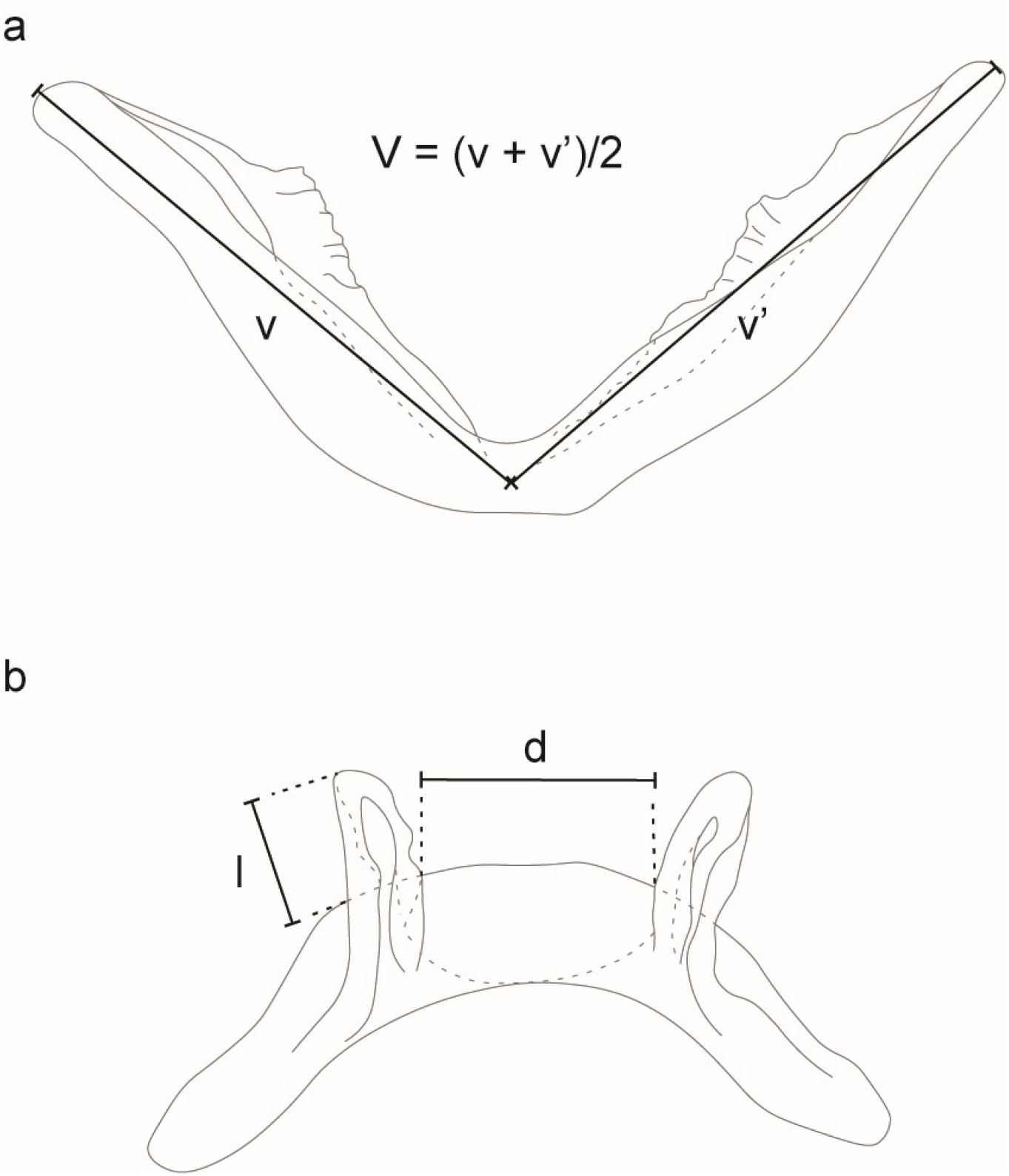
Additional measurements taken on **a** the ventral and **b** the dorsal transverse bar. Abbreviations: V, branch length ; l, auricle length; d, distance between auricles; all as in Douëllou (1993)

The diagnosis-based version of the phylogenetic species concept was adopted for the identification of the new species (Davis & Nixon, 1992). The phylogenetic species concept defines species as reproductively isolated groups of natural populations that originate through a speciation event and end with the next speciation or vanish through extinction (Wägele, 2005). The diagnosis-based version defines a group of specimens as a new species when they consistently differ from another group of specimens in at least one attribute (Davis & Nixon, 1992).

To comply with the regulations set out in article 8.5 of the amended 2012 version of the *International Code of Zoological Nomenclature* (ICZN) (ICZN, 2012), details of the new species have been submitted to ZooBank. The Life Science Identifier (LSID) of the article is XXXX. The Life Science Identifier (LSID) for the new species is reported in the taxonomic summary.

### Morphometric evaluation of interspecific variation

Because the new species closely resembles *C. halli* (see Results), the morphometric variation between specimens of the new species and *C. halli*, collected from *O. niloticus* from Lake Kariba and surrounding fish farms in the present study, was assessed by performing a Principal Component Analysis (PCA) in R version 4.1.0 (R Core Team, 2021). Plots were visualised with the R package *ggplot2* version 3.3.5 (Wickham, 2016). A first PCA was performed including the measurements on both the haptor and MCO. Because the haptor and MCO presumably evolve at a different evolutionary rate (Pouyaud et al., 2006), two additional PCAs were carried out, one including the measurements on the haptor, the other including measurements on the MCO.

### DNA extraction, amplification, sequencing, and alignment

In the genetic analyses, we focused on fragments of the mitochondrial cytochrome *c* oxidase subunit I (*COI*) gene and the small subunit ribosomal DNA (*18S*) and internal transcribed spacer (ITS1) (later referred to as 18S-ITS1). For both gene fragments, specimens of *C. chloeae* sp. nov. infecting *O*. cf. *mortimeri* and *O. niloticus* were selected, as well as specimens of *C. halli* infecting *O. niloticus* and *C. rendalli* (**Table S1**). Micrographs were taken from the sclerotised parts (MCO and haptor) with a Leica DM2500 microscope and the Leica Application Suite X (LASX) software, and deposited as photo vouchers on MorphoBank under the accession numbers XXXX. For DNA extraction, a modified salting-out protocol was followed (provided to us by C. Laumer). Specimens were digested by incubating them in a solution of TNES buffer (400 mM NaCl, 20 mM EDTA, 50 mM Tris pH 8, 0.5% SDS) and 20 mg/mL proteinase *K* at 55°C for one hour. DNA was precipitated by adding 5 M NaCl, 96% (v/v) ethanol, and yeast tRNA as a carrier, and subsequent stored at -20°C for at least one hour. The resulting pellet was purified by two rounds of centrifugation, removing the supernatant and washing the pellet with 70% (v/v) chilled ethanol. The extracted DNA was eluted in 30 µl of 0.1x TE buffer with 0.02% Tween-20 and stored at -20 °C. Amplification was done by a Polymerase Chain Reaction (PCR) with a T100 thermal cycler (Bio-Rad) and BigDye™ Terminator v3.1 Cycle Sequencing Kit (Applied Biosystems). Part of the *COI* gene was amplified and sequenced using the primer pair ASmit1 (5’-TTTTTTGGGCATCCTGAGGTTTAT-3’) and ASmit2 (5’-TAAAGAAAGAACATAATGAAAATG-3’) (Littlewood et al., 1997). The PCR was performed in a reaction mix of 2.5 µL of 10x PCR buffer (Invitrogen), 1 µL of 50 mM MgCl_2_ (Invitrogen), 0.5 µL of 10 mM dNTP mix (Biolegio), 2 µL of each primer (10 µM) (Biolegio), 0.2 µL of 5 U/µL Platinum™ *Taq* Polymerase (Invitrogen), 1 µL template DNA, and 15.8 µL Ultrapure™ DNAse/RNase-free distilled water (Invitrogen) to reach a total volume of 25 µL per reaction. Amplification was carried out under the following conditions: initial denaturation at 95°C for 5 min, 40 cycles of denaturation at 94°C for 1 min, annealing for 1 min at 50°C, and elongation at 72°C for 1 min, and a final elongation at 72°C for 7 min. A nested PCR was performed when the yield of the amplicon was low with a first amplification round using the primer pair ASmit1 and Schisto_3 (5’-TCTTTRGATCATAAGCG-3’) (Lockyer et al., 2003), following the same reaction mix concentrations and PCR conditions as described above, except for the annealing step, which was performed at 44°C. The second amplification round was performed by using the primer pair ASmit 1 and ASmit as described above, using the amplicon of the first round as template DNA.

The 18S-ITS1 fragment was amplified and sequenced using the primer pair S1 (5- ATTCCGATAACGAACGAGACT-3) (Matějusová et al., 2001) and IR8 (5- GCAGCTGCGTTCTTCATCGA-3) (Šimková et al., 2003). The reaction mix contained 3 µL of 10x PCR buffer, 0.9 µL of 50 mM MgCl_2_, 0.6 µL of 10 mM dNTP mix, 1.5 µL of each primer (10 µM) (Biolegio), 0.2 µL of 5 U/µl Platinum™ *Taq* Polymerase, 5 µl template DNA, and 17.3 µL Ultrapure™ DNAse/RNase-free distilled water to reach a total volume of 30 µL per reaction. The PCR was performed under the following conditions: initial denaturation at 94°C for 2 min, 40 cycles of denaturation at 94°C for 1 min, annealing for 1 min at 53°C, and elongation at 72°C for 1.5 min, and a final elongation at 72°C for 10 min. Amplicons were sequenced by Macrogen with bi-directional Sanger sequencing using a 3730xl DNA Analyzer, and the chromatograms of the resulting sequences were checked in Geneious Prime version 2021.2.2 for ends with low base call quality ends, which were manually trimmed. MUSCLE version 3.8.425 (Edgar, 2004) was used under default conditions to align forward and reverse reads, and the consensus sequence was extracted.

### Sequence analyses

To investigate the phylogenetic position of *C. chloeae* sp. nov. within the *Cichlidogyrus-Scutogyrus* monophylum, samples were supplemented with sequences of *C. halli* and other species of *Cichlidogyrus* and *Scutogyrus* from Jorissen et al. (2021), Cruz-Laufer et al. (2021b), and GenBank (**Table S1**).

For both gene fragments, sequences were aligned with MUSCLE under default conditions and overhanging ends were manually trimmed in Geneious Prime. For each gene fragment, the optimal molecular evolution model (GTR+I+G for both gene fragments) was selected based on the corrected Akaike Information Criterion (AICc) using jModelTest2 on the Cipres Science Gateway version 3.3 (Miller et al., 2010). A Bayesian phylogenetic tree was constructed in BEAST version 1.10.4 (Suchard et al., 2018) using a Markov Chain Monte Carlo (MCMC) approach with the best fitting substitution model, a constant size coalescent tree prior (default), and a strict molecular clock model (default). All other operators and prior distributions were left at default settings. Five independent runs were performed from a random starting tree with one cold and one heated chain (deltaTemperature = 0.1) for 10000000 generations with a sampling frequency of 1000. The resulting log files were combined in Tracer version 1.7.2 (Rambaut et al., 2018) with a 50% burn-in to check for convergence in the trace plots. Tree files were combined with LogCombiner (implemented in BEAST) with a 50% burn-in. A Maximum Clade Credibility tree was inferred with default settings in TreeAnnotator (also implemented in BEAST). In addition, a Maximum Likelihood (ML) search was performed in MEGAX version 10.2.6 (Stecher et al., 2020) with 1000 bootstrap replicates using an extensive Subtree-Pruning-Regrafting (SPR level 5) method. Phylogenetic trees were visualised in FigTree version 1.4.4 (Rambaut, 2018). *Cichlidogyrus pouyaudi* Pariselle & Euzet, 1994 was used as an outgroup to root the phylogenetic trees based on the 18S-ITS1 fragment because of the basal position of this species in the phylogenetic tree of the *Cichlidogyrus*-*Scutogyrus* monophylum (Mendlová et al., 2012; Messu Mandeng et al., 2015). The *COI* fragment of *C. pouyaudi* is not available on GenBank. Therefore, the phylogenetic tree based on this gene fragment was rooted with *Cichlidogyrus falcifer* Dossou & Birgi, 1984, the *COI* fragment of which is available, because this species falls in a different clade than specimens of *C. halli* and *C. chloeae* sp. nov. in the phylogenetic tree based on the 18S-ITS1 fragment (see Results).

The intraspecific genetic distances between specimens of *C. chloeae* sp. nov. and the interspecific genetic distances between specimens of *C. halli* and *C. chloeae* sp. nov. were calculated in MEGAX using the Kimura-2-parameter (K2P) distance model (Kimura, 1980) based on both the *COI* and 18S-ITS1 fragment, supplementing our dataset with GenBank sequences where available (**Table S1**).

## Results

A total of 27 specimens of *O*. cf. *mortimeri* and 29 specimens of *C. rendalli* were caught in Lake Kariba. Additionally, 58 specimens of *O. niloticus* were collected: 27 from aquaculture facilities (Lake Harvest and Nicholson Bream Farm) and 31 from Lake Kariba. On *O*. cf. *mortimeri*, a total of 63 specimens of *C. chloeae* sp. nov. was found, while no specimens of *C. halli* were detected. On *C. rendalli*, three specimens of *C. chloeae* sp. nov. and one specimen of *C. halli* were found. On *O. niloticus*, a total of 40 specimens of *C. chloeae* sp. nov. was found on feral fish from the lake, while none were found on farmed fish. A total of 16 specimens of *C. halli* were found on feral *O. niloticus*, and 203 specimens on farmed fish. An overview of these results is given in **Table 2**. Apart from specimens of *C. halli* and *C. chloeae* sp. nov., also specimens belonging to other species of Monogenea were found on the gills of these hosts (for detailed information see Geraerts et al. (In review)).

**Table 2.** Number of specimens of *C. halli* and *C. chloeae* sp. nov. found on the fish species studied; number of fish specimens (n) between brackets.

| Parasite species | *O. cf. mortimeri* (n = 27) | *Coptodon rendalli* (Boulenger, 1897) (n = 29) | *Oreochromis niloticus* (Linnaeus, 1758) (n = 58) |  |
| --- | --- | --- | --- | --- |
|  |  |  | Farmed (n = 27) | Feral (n = 31) |
| *C. chloae* sp. nov | 63 | 3 | / | 40 |
| *C. halli* | / | 1 | 203 | 16 |

The species description of *C. chloeae* n.sp. is presented below, together with a morphological comparison with *C. halli*. For the measurements on the hard parts, 10 specimens of *C. chloeae* sp. nov. from *O*. cf. *mortimeri*, 18 specimens from *O. niloticus*, and three specimens from *C. rendalli* were at our disposal. Additionally, measurements were taken from 21 specimens of *C. halli* from *O. niloticus* (20 specimens from farmed hosts and one from a feral host) that were available for morphological analyses. Measurements on both species can be found in **Table 3**.

**Table 3.**
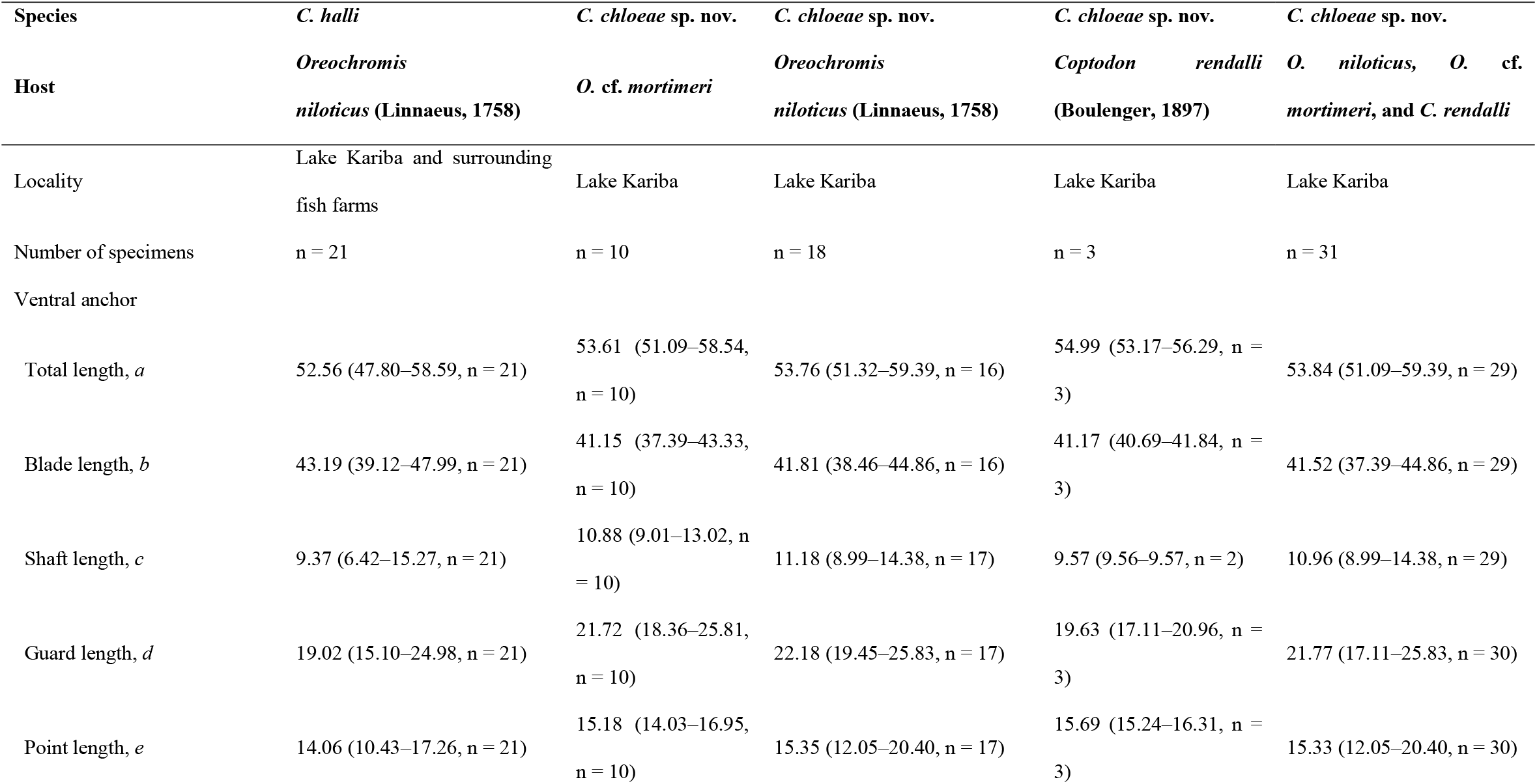
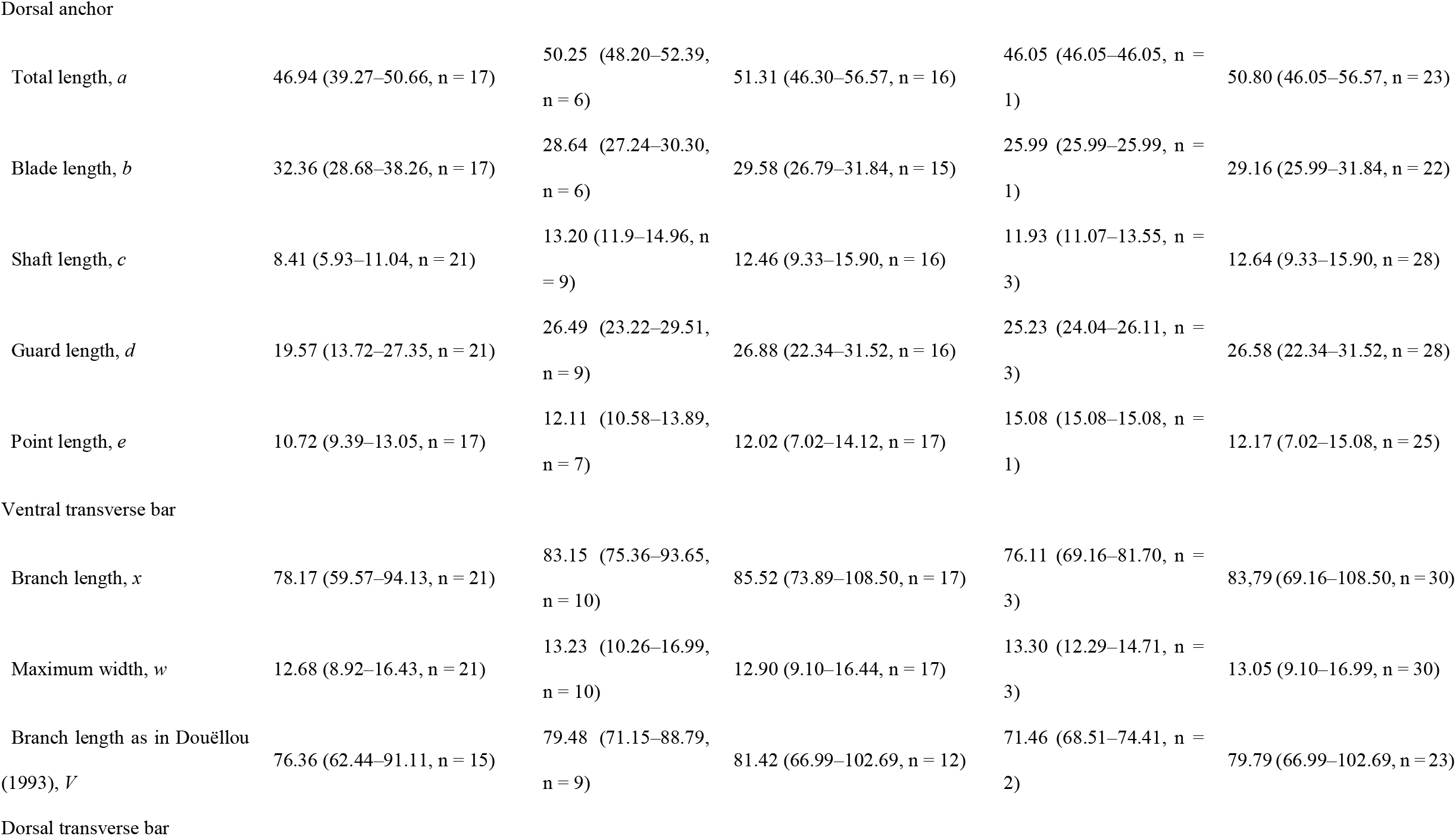
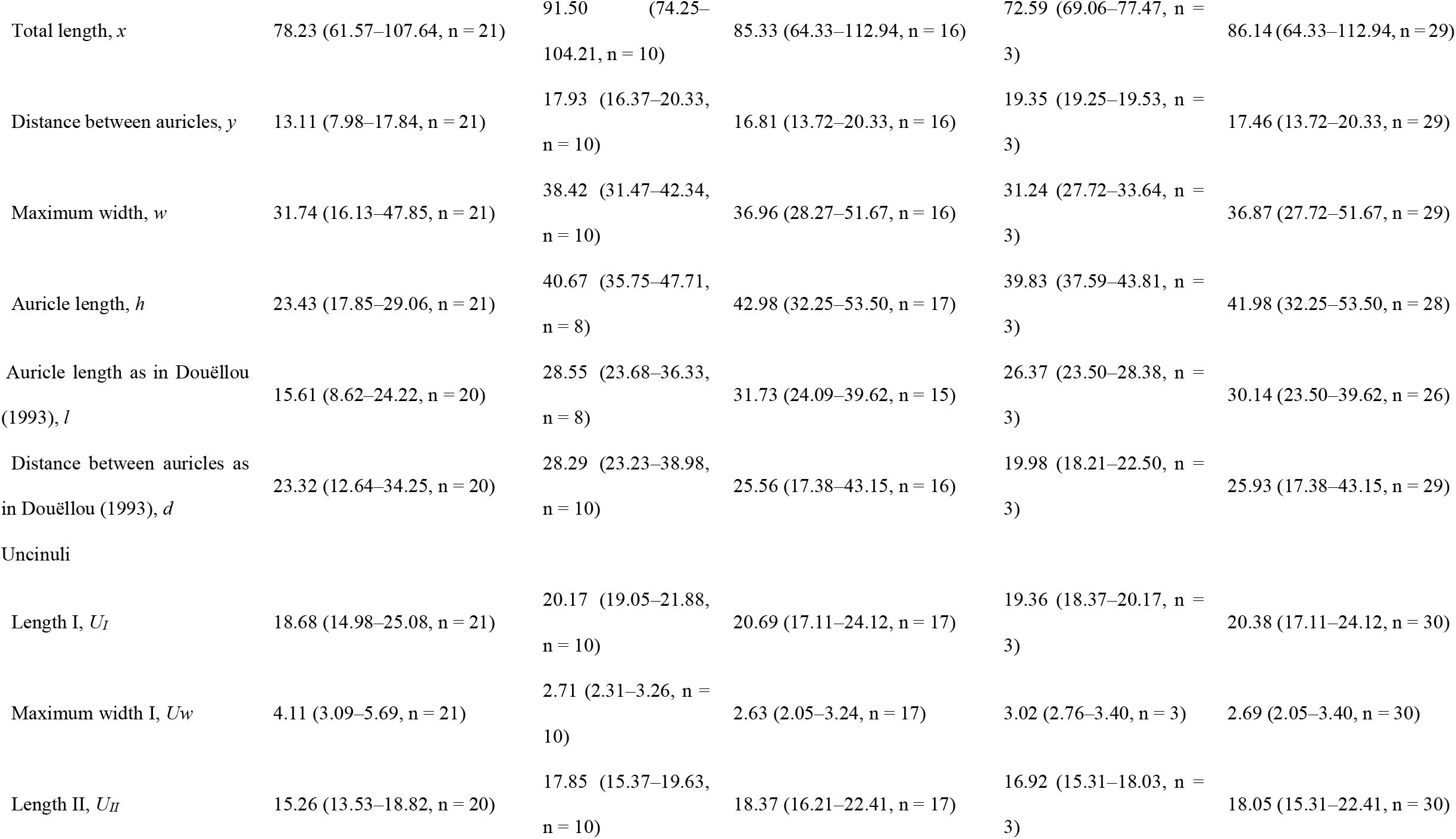
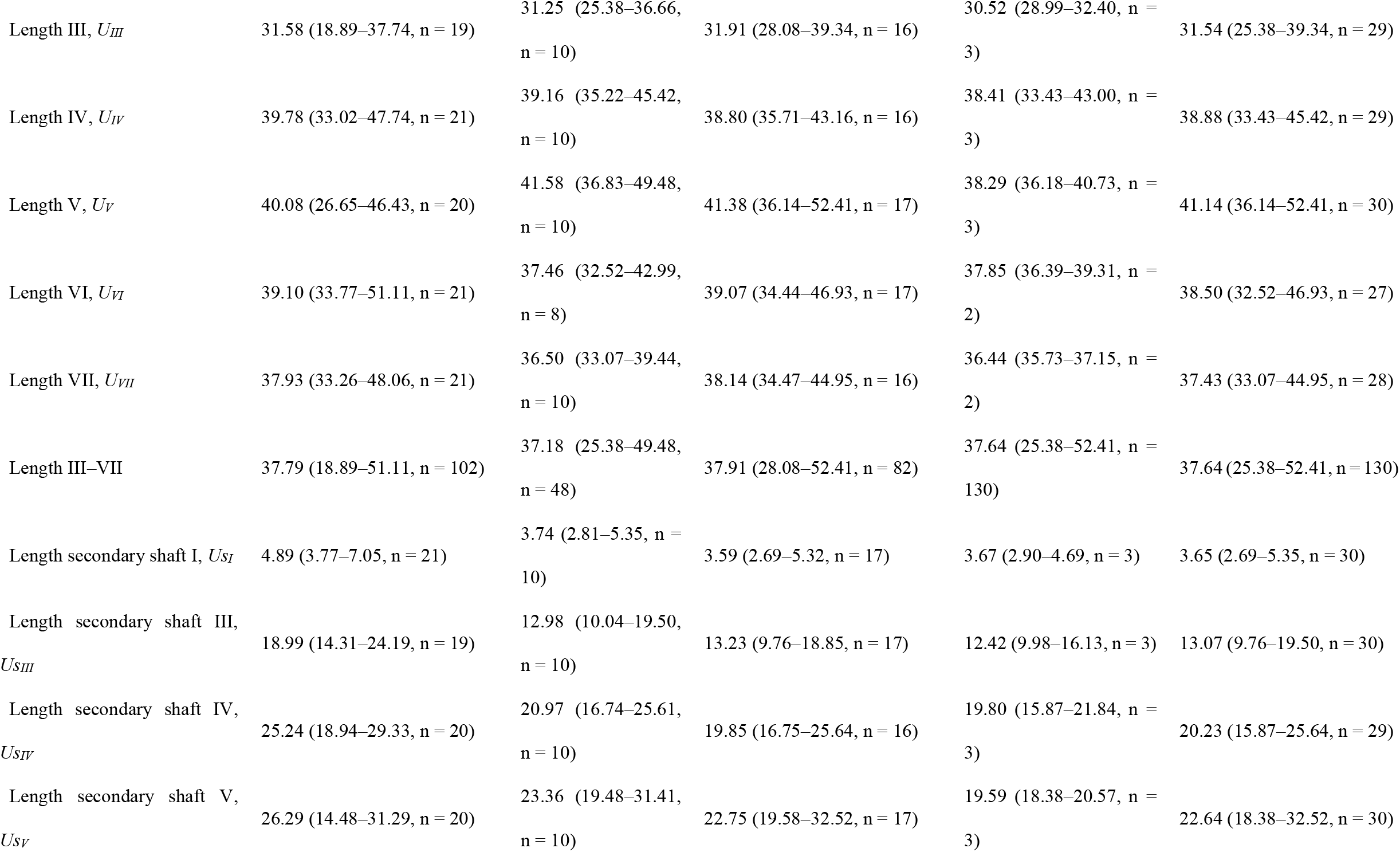
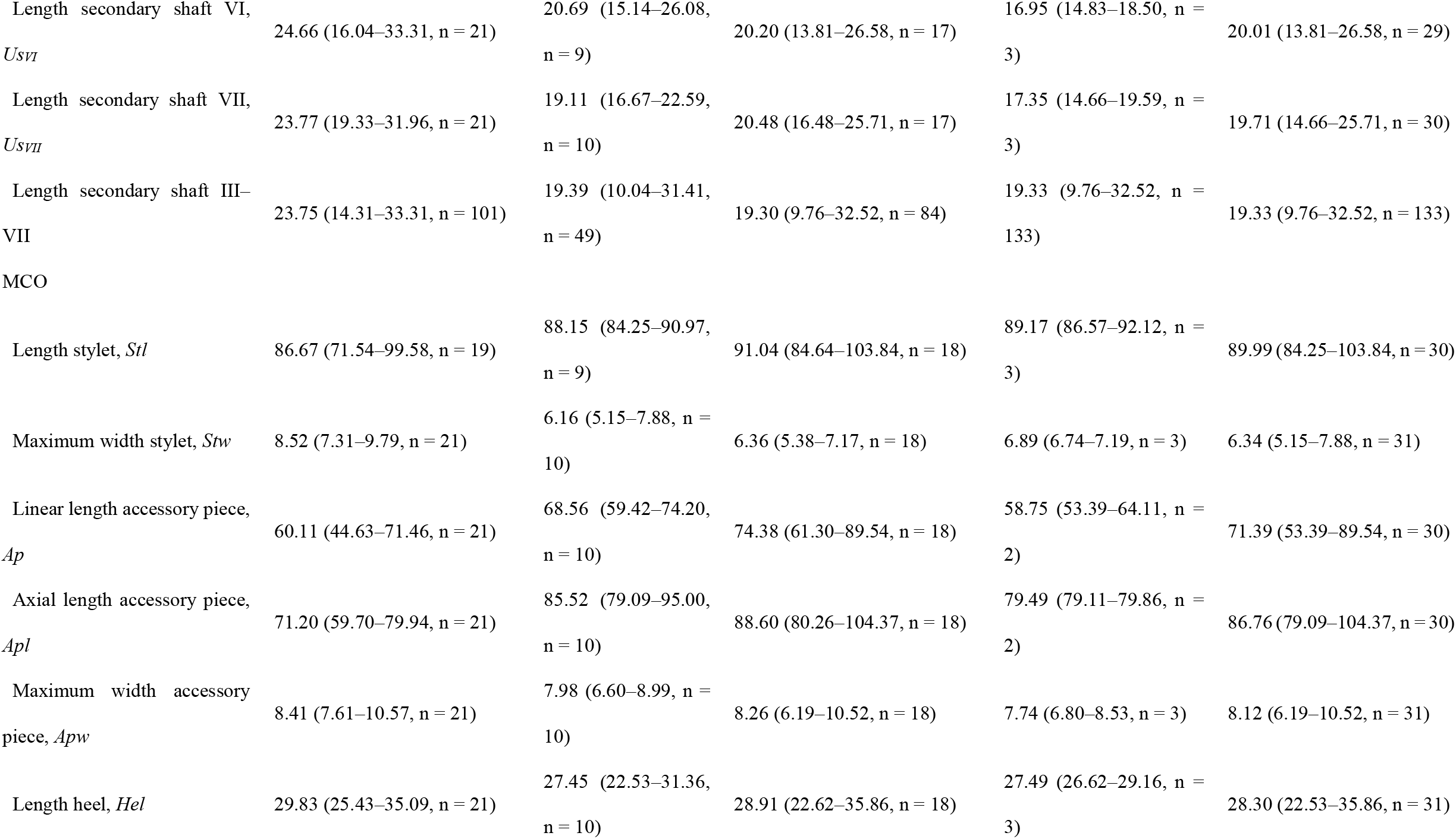
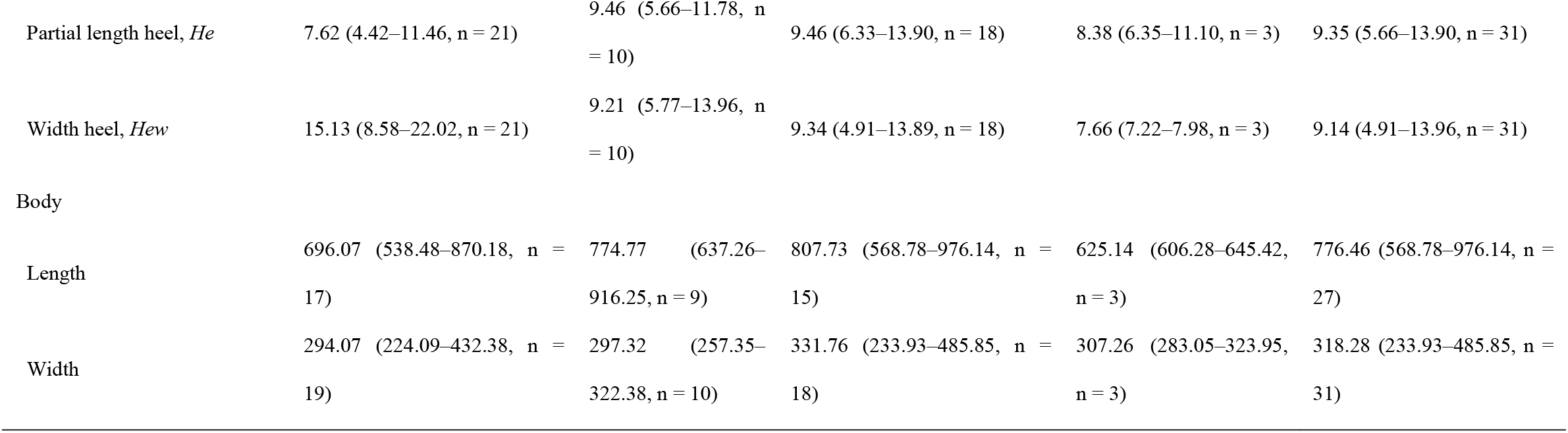
Measurements (in μm) on *C. halli* and *C. chloeae* sp. nov. Note: Measurements are given as the mean followed by the range and number of measured specimens (n) in parentheses.

### Taxonomy

Family Dactylogyridae Bychowski, 1933

Genus *Cichlidogyrus* Paperna, 1960

*Cichlidogyrus chloeae* Geraerts sp. nov.

HOLOTYPE: KN.28605.

PARATYPES: KN.28606–KN.28613, RMCA_VERMES_43649– RMCA_VERMES_43658, and UH nos. XXXX.

TYPE LOCALITY: Lake Kariba, Zimbabwe.

HABITAT: Gills of host

TYPE HOST: *Oreochromis* cf. *mortimeri* (Perciformes: Cichlidae).

OTHER HOSTS: *Oreochromis niloticus* (Linnaeus 1758) and *Coptodon rendalli* (Boulenger, 1897) (Perciformes: Cichlidae).

PREVALENCE AND INTENSITY: see **Table 4**.

**Table 4.**
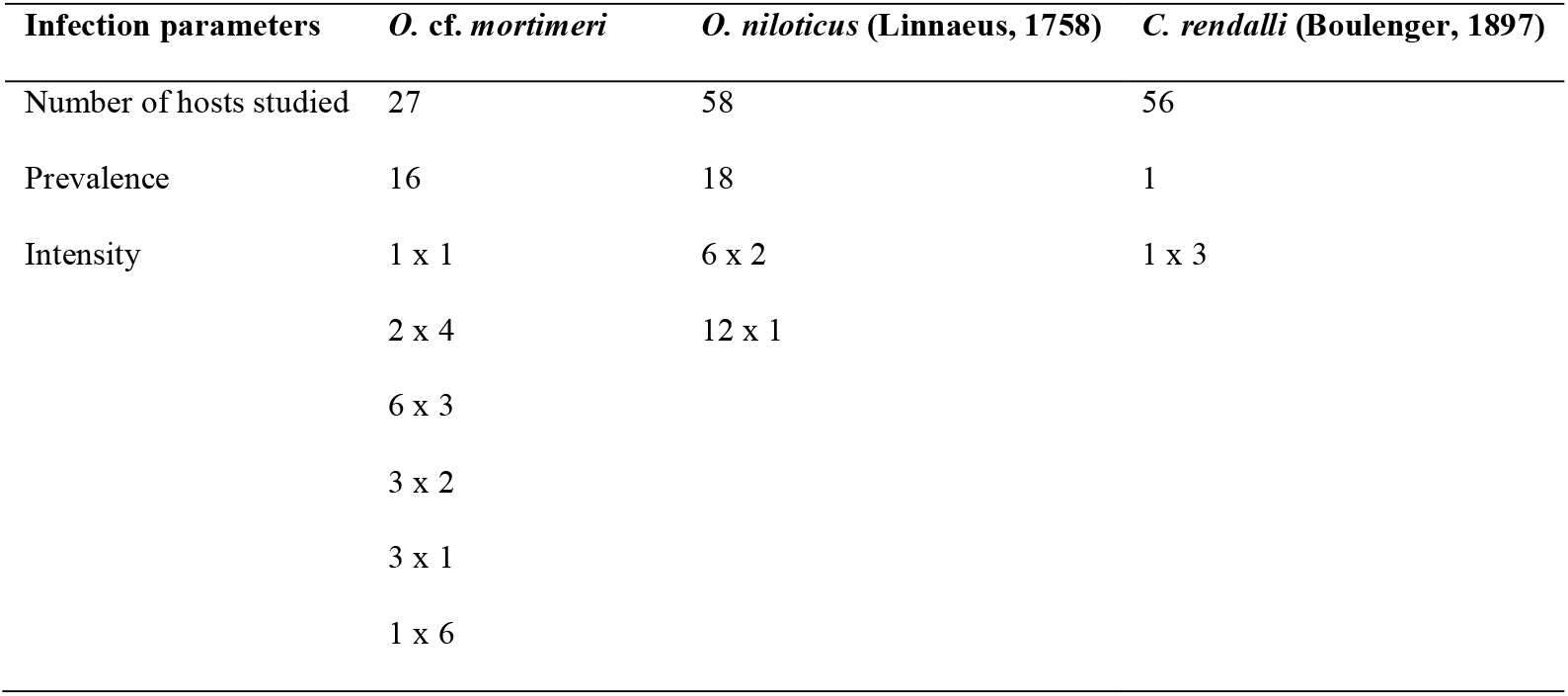
Prevalence and intensity as defined by Bush et al. (1997) of *C. chloeae* sp. nov. on the studied hosts with the intensity expressed as ‘number of host specimens x number of parasites infecting these hosts’.

ZOOBANK REGISTRATION: The Life Science Identifier (LSID) for *Cichlidogyrus chloeae* sp. nov. is XXXX.

ETYMOLOGY Dedicated to the first author’s best friend and support Chloë Vervoort. DIAGNOSIS: Species of *Cichlidogyrus* with small uncinuli I (length ±20 μm) and long uncinuli III to VII (length ±38 μm), large ventral anchors (total length ±54 μm) with an asymmetrical base (guard length ±22 μm, shaft length ±11 μm) and large dorsal anchors (total length ±51 μm) with an asymmetrical base (guard length ±27 μm, shaft length ±13 μm). Dorsal transverse bar with large auricles (length ±42 μm) and ventral bar with distinct wing-shaped attachments. Penis stylet tubular (length ±90 μm) and broad (maximum width ±6 μm) and accessory piece (axial length ±87 μm, maximum width ±8 μm) with a triangular shaped cap at distal end.

### Description (Fig. 3; Fig. 4a–b)

**Fig. 3.**
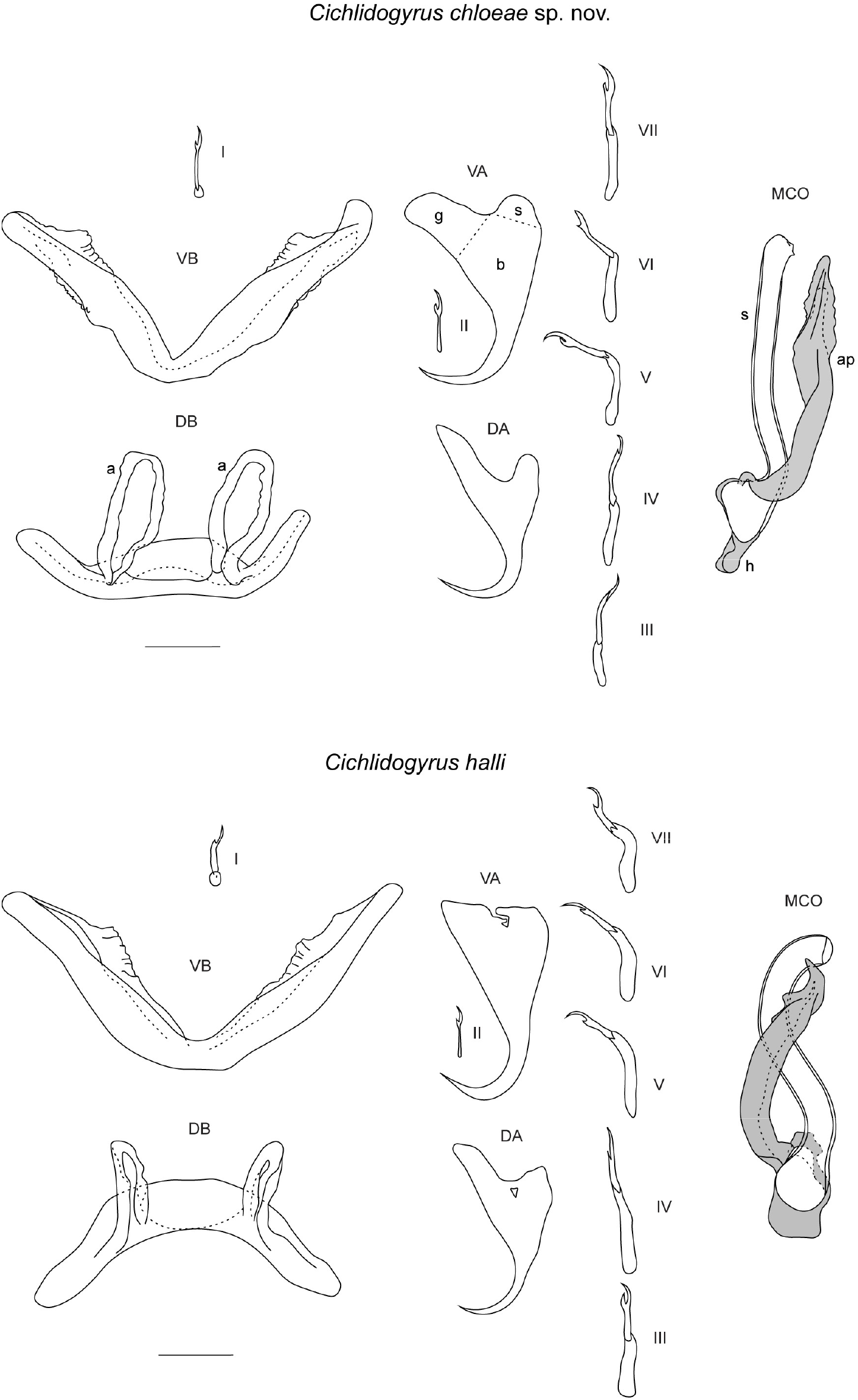
Drawings of the sclerotised structures of *Cichlidogyrus chloeae* sp. nov. (top) and *C. halli* (bottom). Drawings of *C. chloeae* sp. nov. are based on two specimens: the holotype KN.28605 for the MCO, and paratype XXXX for the haptor. Drawings of *C. halli* are based on three specimens: voucher XXXX for the ventral bar and anchors, voucher XXXX for the dorsal bar, and voucher XXXX for the uncinuli and MCO. Abbreviations: I–VII, uncinuli; VA, ventral anchor (g, guard; s, shaft; b, blade); VB, ventral transverse bar; DA, dorsal anchor; DB, dorsal transverse bar (a, auricle); MCO, male copulatory organ with penis stylet (s) in white, and accessory piece (ap) and heel (h) in grey. Scale bar: 20 µm.

**Fig. 4.**
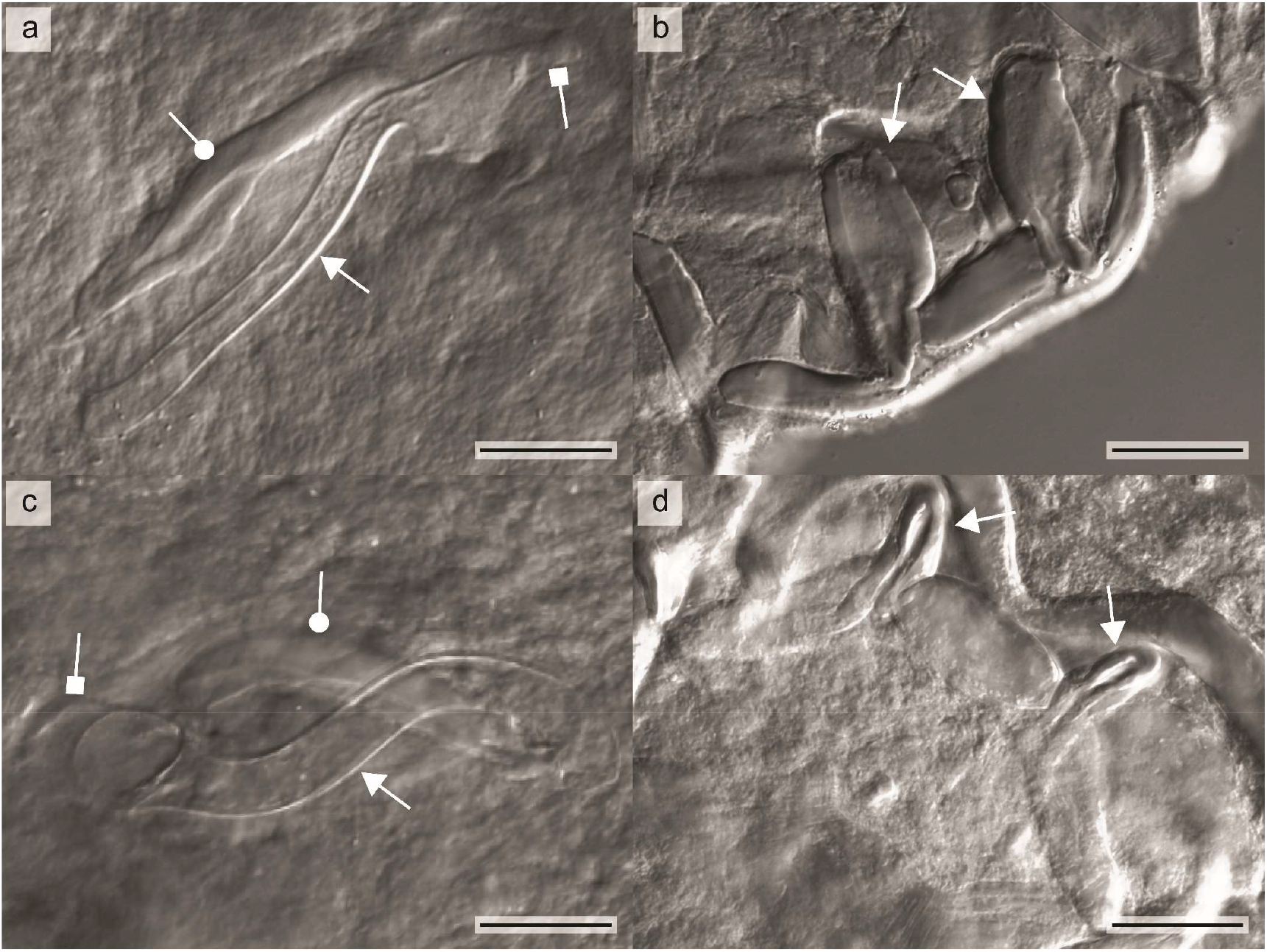
Micrographs of the MCO and dorsal transverse bar of *C. chloeae* sp. nov. and *C. halli*. **a** MCO and **b** dorsal transverse bar of *C. chloeae* sp. nov. **c** MCO and **d** dorsal transverse bar of *C. halli*. The arrows in **a** and **c** indicate the penis stylet, the circle the accessory piece, and the square the heel. Arrows in **b** and **d** indicate the auricles of the dorsal bar. Scale bar: 20 µm.

[Based on 31 specimens; metrical data in **Table 3**]

#### HAPTOR

Anchors 2 pairs. Ventral anchors large with massive asymmetrical base; guard and shaft broad with guard approximately 2 times as long as shaft. Dorsal anchors of about same total length as ventral anchors; base asymmetrical with guard and shaft narrower than those of ventral anchor; guard approximately 2 times as long as shaft. Blades of both ventral and dorsal anchors arched. Ventral transverse bar V-shaped, with 2 long branches with distinct wing-shaped attachments along distal half. Dorsal transverse bar large and made up of thick midsection, tapering towards its extremities, and 2 pronounced auricles inserted at its dorsal face. Uncinuli 7 pairs; uncinuli I short with small round secondary shaft; uncinuli III to VII long; uncinuli III on average shorter than uncinuli IV–VII (Uncinuli are called short or long based on their standardised length i.e. the division of their total length by the total length of the second uncinuli, which retain their larval size (Pariselle & Euzet, 2009)).

#### MALE GENITALIA

MCO consisting of a long penis stylet, accessory piece and heel. Penis stylet broad and tubular with constant width along its length and an enlarged proximal irregularly shaped basal bulb. Accessory piece about the same axial length as penis stylet and proximally connected to base of stylet; broader than penis stylet with an elongated triangular shaped cap at distal end. Heel pronounced and attached to base of penis stylet, narrower than basal bulb of penis stylet.

#### FEMALE GENITALIA

No sclerotised vagina visible.

### Morphological discussion and morphometric evaluation of interspecific variation between *C. chloeae* sp. nov. and *C. halli*

The described species is classified as a species of *Cichlidogyrus* because it shows all diagnostic features of the genus: two pairs of anchors (one dorsal and one ventral), two transverse bars (ventral transverse bar V-shaped, dorsal transverse bar with two auricles, 14 uncinuli, and an MCO consisting of a penis stylet and an accessory piece (Paperna, 1960; Pariselle & Euzet, 2009). It belongs to the group of species of *Cichlidogyrus* with small uncinuli I and long uncinuli III to VII (Pariselle & Euzet, 2009). Based on the morphology of the sclerotised structures, *C. chloeae* sp. nov. resembles *C. halli*: both species have small uncinuli I and long uncinuli III to VII, large anchors with an asymmetrical base, a broad tubular stylet and a triangular shaped cap at the distal end of the accessory piece. However, *C. chloeae* sp. nov. differs from *C. halli* in the length of the auricles, which are almost twice as long in *C. chloeae* sp. nov. Also, the penis stylet is slightly wider, the accessory piece longer with more elongated and less pronounced triangular cap, and the heel narrower in *C. chloeae* sp. nov. compared to *C. halli*, in which the heel engulfs the entire basal bulb of the penis stylet (**Fig. 3**; **Fig. 4**).

In the PCA including measurements on both haptor and MCO, the first three principal components explain respectively 42.7%, 21.1% and 9.7% of the variation (**Fig. 5a**; **Fig. S1a**). The linear (*Ap*) and axial length of the accessory piece (*Apl*), and the length of the auricles of the dorsal transverse bar (*DBh*) have the highest contribution to PC1 as well as to PC2. The first three principal components of the PCA including only measurements on the haptor explain respectively 39.5%, 22.9% and 8.7% of the variation (**Fig. 5b**; **Fig. S1b**). The total length of the dorsal transverse bar (*DBx*), the length of the auricles of the dorsal transverse bar (*DBh*) and the length of the branches of the ventral bar (*VBx*) contribute most to PC1, while the total length of the dorsal transverse bar (*DBx*), the length of the auricles of the dorsal transverse bar (*DBh*) and the length of the secondary shaft of uncinuli III (*IIIus*) contribute most to PC2. Finally, in the PCA including only measurements on the MCO, the first three principal components explain 73.8%, 10.3%, and 7.6% of the variation, respectively (**Fig. 5c**; **Fig. S1c**). In this PCA, the linear (*Ap*) and axial length (*Apl*) of the accessory piece, and the length of the penis stylet contribute most to both PC1 and PC2. Each biplot shows two clusters: one including specimens of *C. chloeae* sp. nov., the other including specimens of *C. halli*. The measurement contributing to this clustering (i.e. pointing in the direction perpendicular to the clusters) is the length of the auricles of the dorsal transverse bar (DBh) and the length of the accessory piece (**Fig. 5**).

**Fig. 5.**
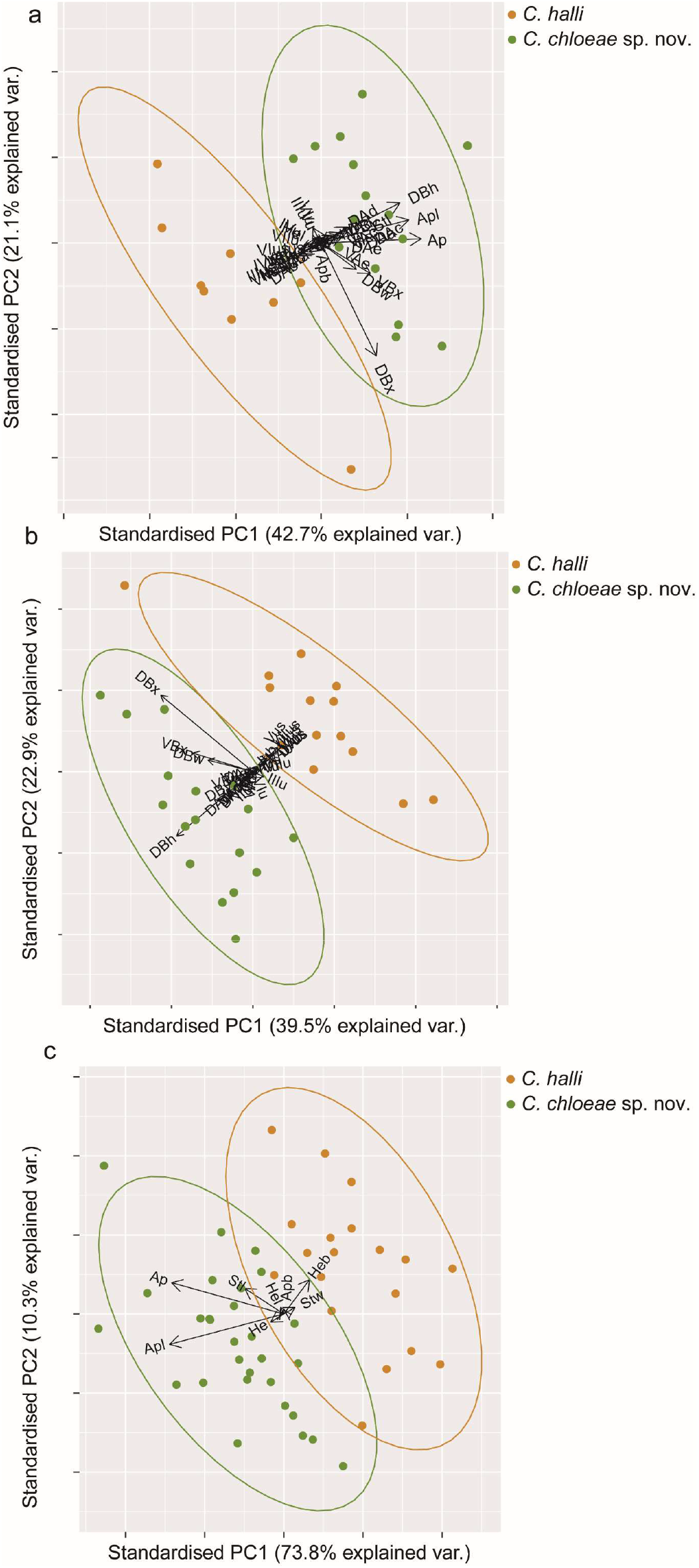
Biplots of the PCAs plotting the first two principal components PC1 and PC2: **a** PCA based on all measurements, **b** measurements on the haptor only, and **c** measurements on the MCO only. Each dot represents one specimen. Different colours represent different species i.e. *C. halli* and *C. chloeae* **sp. nov**. Ellipses are drawn at a confidence interval of 0.95. The contribution of the different measurements to the principal components are shown by arrows.

### Phylogenetic position of *C. chloeae* sp. nov. within the *Cichlidogyrus*-*Scutogyrus* monophylum

In the Bayesian phylogenetic tree inferred from the 18S-ITS1 fragment, specimens of *C. chloeae* sp. nov. fall in a well-supported monophyletic clade together with specimens of *C. halli*, with the ‘*C. chloeae* sp. nov.’ clade being the sister group of the ‘*C. halli’* clade (**Fig. 6**). The same sister group relationship is suggested by the topology of the ML phylogenetic tree inferred from the same gene fragment albeit with weaker support (bootstrap value <0.85) (**Fig. S2**).

**Fig. 6.**
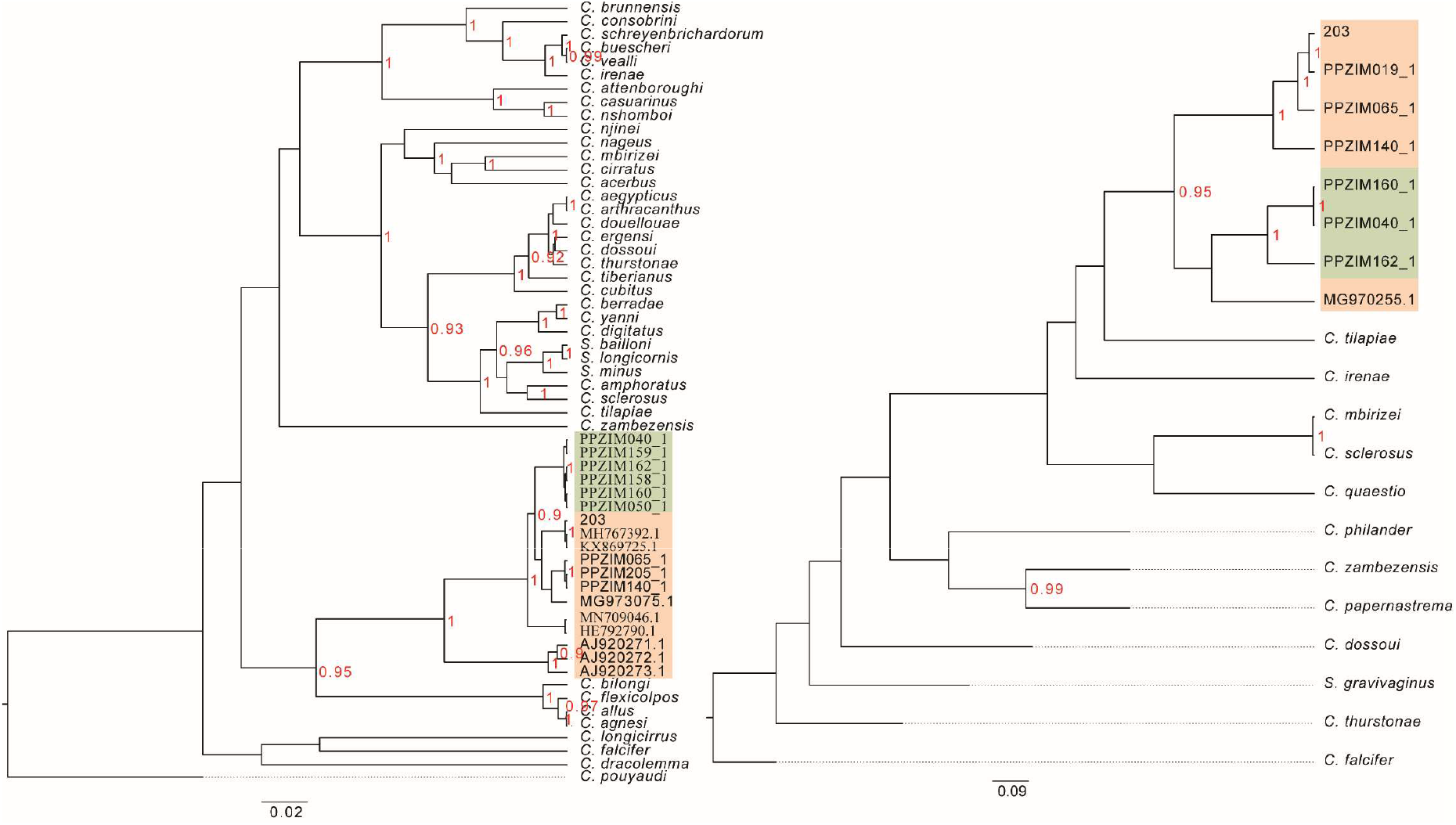
Bayesian phylogenetic trees of specimens of *Cichlidogyrus* and *Scutogyrus* inferred from the 18S-ITS1 fragment (left) and *COI* fragment (right). Only well supported nodes (bootstrap values ≥ 0.85) are indicated by support values (in red). Scale bar indicates number of substitutions per site. Specimens of *C. chloeae* sp. nov. framed in green, specimens of *C. halli* framed in orange. Parasite labels and GenBank accession numbers of the included specimens can be found in **Table S1**.

Also, in the Bayesian (**Fig. 6**) and ML phylogenetic tree (**Fig. S2**) inferred from the *COI* fragment, specimens of *C. chloeae* sp. nov. fall in the same clade as specimens of *C. halli* with one specimen of *C. halli* (MG970255.1) forming the sister group of the clade including the specimens of *C. chloeae* sp. nov., though with low support values (bootstrap value <0.85).

The barcoding gap between the intraspecific genetic distances of specimens of *C. chloeae* sp. nov. and the intraspecific distances between specimens of *C. chloeae* sp. nov. and *C. halli* ranged from 0.131 to 0.161 based on the *COI* fragment, and from 0 to 0.028 based on the 18S-ITS1 fragment.

## Discussion

In the present study, we describe *C. chloeae* sp. nov. and make a morphological and morphometric comparison of *C. chloeae* sp. nov. with *C. halli*. Based on the measurements and drawings, *C. chloeae* sp. nov. can easily be distinguished from *C. halli* by longer auricles of the dorsal transverse bar and a longer accessory piece, a slightly wider penis stylet, a narrower heel, and a triangular cap of the accessory piece being more elongated and less pronounced in *C. chloeae* sp. nov. Therefore, it is clear that *C. chloeae* sp. nov. is a new species.

*C. chloeae* sp. nov. and *C. halli* are closely related, falling in the same monophyletic clade based on both the nuclear and mitochondrial gene fragment. This finding adds to the growing evidence of the presence of a ‘*C. halli* complex’, encompassing several (undescribed) species as proposed by Jorissen et al. (2018b), Jorissen et al. (2021) and Geraerts et al. (In press). A barcoding gap between 13 and 16% is found based on the *COI* fragment which is in accordance to the findings of Jorissen et al. (2021) (barcoding gap at 15% for the *COI* gene) and Geraerts et al. (In Press). The gap we found between the intra- and interspecific genetic distances based on the 18S-ITS1 fragment is much smaller (between 0 and 3%), which is consistent with the findings of Jorissen et al. (2021) and Geraerts et al. (In review).

### Previous mix-up of C. chloeae sp. nov. with C. halli?

The sole study collecting gill parasites of cichlids in Lake Kariba was carried out in the 1990s (Douëllou, 1993). The species of *Cichlidogyrus* found on *O. mortimeri* were *C. halli*, C. *dossoui* Douëllou, 1993, *C. karibae* Douëllou, 1993, *C. tilapiae* Paperna, 1960, *C. sclerosus* Paperna & Thurston, 1969, and *C. zambezensis* Douëllou, 1993. The species of *Cichlidogyrus* found on *C. rendalli* were *C. dossoui, C. quaestio* Douëllou, 1993, and *C. tiberianus* Paperna, 1960. In Douëllou’s research, the morphometrics of specimens of *C. halli* were compared with those made by Price & Kirk (1967) in the original species description. She already reported a difference in the auricle length between the specimens of *C. halli* found in her study and those from the original description (Douëllou, 1993), but did not recognise it as a new species. In the present study, the auricle length of *C. chloeae* sp. nov. overlaps with ‘*C. halli*’ found in the study of Douëllou (1993), while the auricle length of *C. halli* found in our study overlaps with that of *C. halli* described in the original species description by Price & Kirk (1967) (**Table 5**). We are, therefore, convinced of the fact that the specimens of ‘*C. halli*’, found by Douëllou (1993), are actually specimens of *C. chloeae* sp. nov.

**Table 5.** Measurements (in μm) on *C. halli* from Price & Kirk (1967), Douëllou (1993) and the present study, and measurements on *C. chloeae* sp. nov. from the present study.

| Species | *C. halli* | *C. halli* | *C. halli* | *C. chloae* sp. nov. |
| --- | --- | --- | --- | --- |
| Host | *Oreochromis shiranus* Boulenger, 1897 | *Oreochromis cf. mortimeri* | *Oreochromis niloticus* (Linnaeus, 1758) | *O. niloticus*, *O. mortimeri*, and *C. rendalli* |
| Locality | Shire River, Malawi | Lake Kariba, Zimbabwe | Lake Kariba, Zimbabwe | Lake Kariba, Zimbabwe |
| Number of specimens | n = 8 | n = 15 | n = 21 | n = 31 |
| Reference | Price & Kirk, 1967 | Douëllou, 1993 | This study | This study |
| Ventral anchor |  |  |  |  |
| Total length, *a* | 54–62 | 49–60 | 52.56 (47.80–58.59, n = 21) | 53.84 (51.09–59.39, n = 29) |
| Dorsal anchor |  |  |  |  |
| Total length, *a* | 53–60 | 42–56 | 46.94 (39.27–50.66, n = 17) | 50.80 (46.05–56.57, n = 23) |
| Ventral transverse bar |  |  |  |  |
| Branch length, *V* | 104–122 | 104–144 | 76.36 (62.44–91.11, n = 15) | 79.79 (66.99–102.69, n = 23) |
| Dorsal transverse bar |  |  |  |  |
| Total length, *x* | 68–79 | 51–73 | 78.23 (61.57–107.64, n = 21) | 86.14 (64.33–112.94, n = 29) |
| Auricle length, *l* | 14 | 20–25 | 15.61 (8.62–24.22, n = 20) | 30.14 (23.50–39.62, n = 26) |
| Uncinuli |  |  |  |  |
| Length I, *U<sub>I</sub>* | 20–22 | 17–20 | 18.68 (14.98–25.08, n = 21) | 20.38 (17.11–24.12, n = 30) |
| Length II, $U_{II}$ | 20–22 | 16–18 | 15.26 (13.53–18.82, n = 20) | 18.05 (15.31–22.41, n = 30) |
| Length III–VII | 35–44 | 29–43 | 37.79 (18.89–51.11, n = 102) | 37.64 (25.38–52.41, n = 130) |
| MCO |  |  |  |  |
| Length stylet, $Stl$ | 82–86 | 66–96 | 86.67 (71.54–99.58, n = 19) | 89.99 (84.25–103.84, n = 30) |
| Axial length accessory piece, $Apl$ | 61–67 | 54–66 | 71.20 (59.70–79.94, n = 21) | 86.76 (79.09–104.37, n = 30) |
| Body |  |  |  |  |
| Length | 525–721 | 700–1400 | 696.07 (538.48–870.18, n = 17) | 776.46 (568.78–976.14, n = 27) |
| Width | 160–205 | 220–340 | 294.07 (224.09–432.38, n = 19) | 318.28 (233.93–485.85, n = 31) |

This hypothesis is further supported by the fact that *C. halli* is abundant on farmed *O. niloticus*, but is not found on *O*. cf. *mortimeri* in the present study. Furthermore, *C. chloeae* sp. nov. is found on *O*. cf. *mortimeri* and feral *O. niloticus*, but not on farmed *O. niloticus* (**Table 2**).

### Feral Nile tilapia as reservoir for C. *chloeae* sp. nov

The absence of *C. chloeae* sp. nov. on farmed *O. niloticus*, and its presence on feral *O. niloticus*, suggests a host-switch from *O*. cf. *mortimeri* to *O. niloticus*. The few specimens of *C. chloeae* sp. nov. that were found on *C. rendalli* also suggest a host switch from *O*. cf. *mortimeri* to *C. rendalli* as it was not yet detected on *C. rendalli* by earlier research. Indeed, Douëllou (1993) did not find ‘*C. halli’* (now *C. chloeae* sp. nov.) on *C. rendalli* either.

Invasion ecology often focuses on spillover of introduced parasites to native hosts, though infection of non-indigenous hosts by native parasites can also pose a potential threat to native species. Non-indigenous hosts can act as a new reservoir for native parasites, providing an additional habitat in which the parasite can persist and reproduce. Ultimately, this can lead to an expansion of the parasite population and potentially increase the prevalence and intensity of this parasite on its native host by spillback (Goedknegt et al., 2016; Kelly et al., 2009; Poulin et al., 2011).

Species of *Cichlidogyrus* are regarded as being highly host specific. However, *C. chloeae* sp. nov. is classified as an intermediate generalist (using the terminology proposed by Mendlová & Šimková 2014), infecting non-congeneric cichlids of different tribes, i.e. Oreochromini and Coptodonini. This, together with their one-host life cycle (Řehulková et al., 2018) and the fact that different species of tilapia are morphologically and ecologically similar (Vignon et al., 2011), could facilitate host switches between introduced and native tilapias in Lake Kariba.

## Acknowledgements

We appreciate the help of the people involved in the field work and sampling procedure. Furthermore, we would like to thank Christopher Laumer (EMBL-European Bioinformatics Institute, UK) and Gontran Sonet (Royal Belgian Institute of Natural Sciences, Belgium) for providing us with the extraction protocol and R scripts, respectively. Finally, we would like to thank Ria Vanderspikken (Hasselt University, Belgium) and Natascha Steffanie (Hasselt University, Belgium) for their technical support in the laboratory.

## Figures

**Fig. S1.**
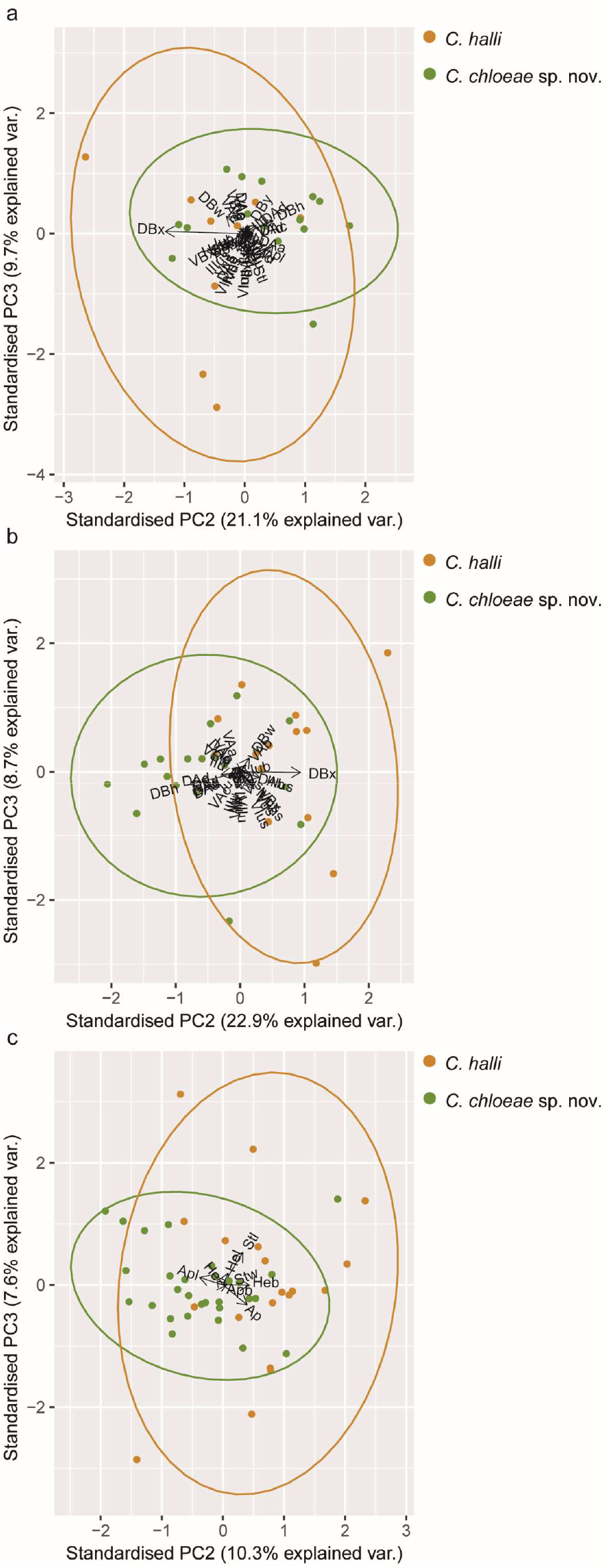

**Fig. S2.**
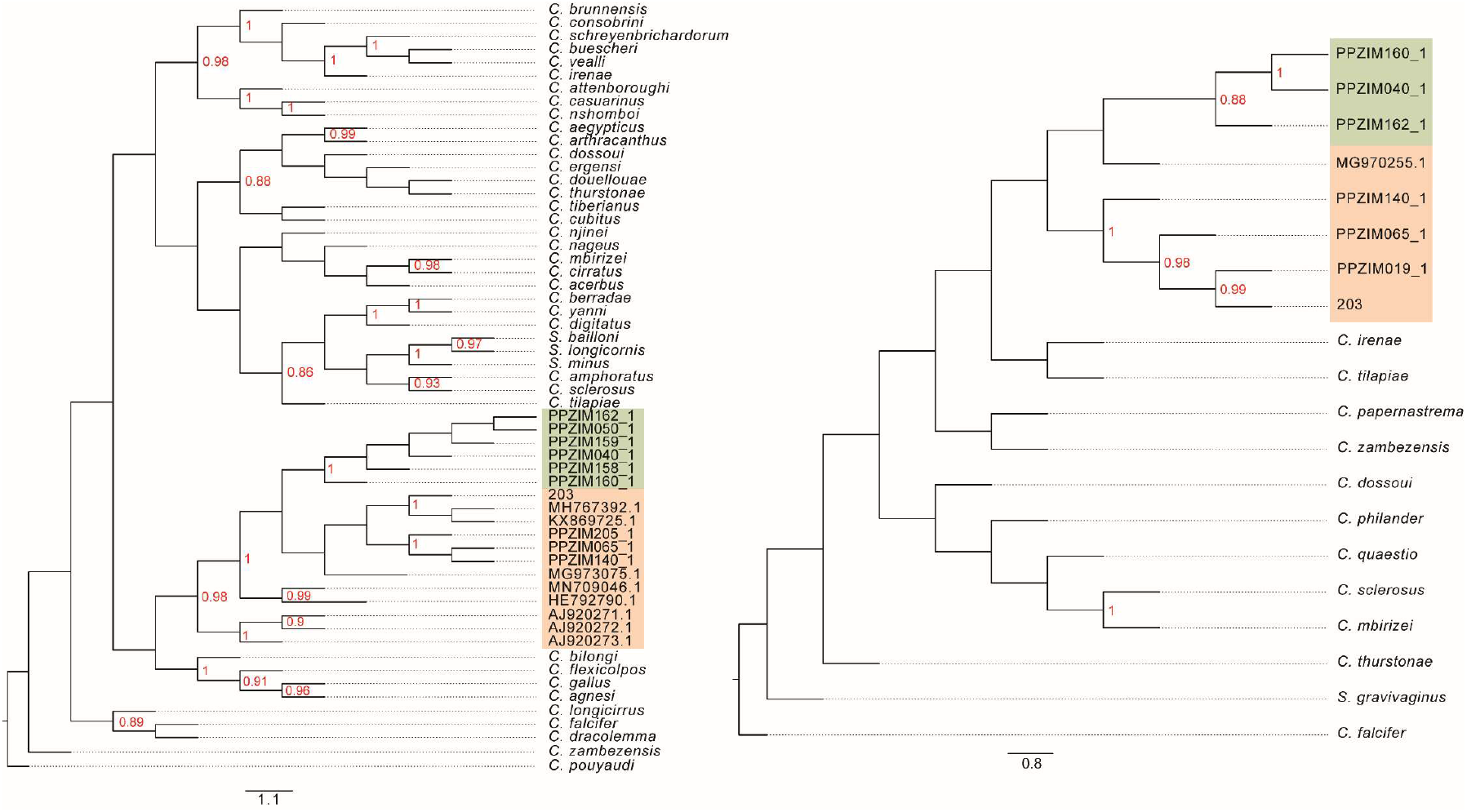

## Footnotes

1 The term ‘tilapia’ will be used in the present study to refer to a paraphyletic group of cichlids consisting of several haplotilapiine tribes, including commercially important genera, such as *Oreochromis* Günther 1889, *Tilapia* Smith 1840, *Coptodon* Gervais 1853, and *Sarotherodon* Rüppell 1852 (Dunz & Schliewen, 2013; Trewavas, 1982).

