## Supplementary material for "A new species of *Cichlidogyrus* Paperna, 1960 (Platyhelminthes: Monogenea: Dactylogyridae) infecting tilapias in Lake Kariba (Zimbabwe), with a discussion on its phylogenetic position"

### Table S1

Specimens of *Cichlidogyrus* Paperna, 1960 and *Scutogyrus* Pariselle & Euzet, 1995 selected for genetic analyses with details about the parasite species, host species, host label, parasite label, GenBank accession number, country and basin of sampling, and location label (L). Location labels correspond to the ones in **Fig. 1**. Parasites with labels starting with ‘PPZIM’ are the specimens collected in the present study.

| **Parasite species** | **Host species** | **Host label** | **Parasite label** | **Genbank accession number** | **Country** | **Basin** | **L** |
| --- | --- | --- | --- | --- | --- | --- | --- |
| **18S-ITS1** |  |  |  |  |  |  |  |
| *C. chloeae* **sp. nov.** | *Oreochromis* cf. *mortimeri* | ZIM046 | PPZIM158.1 | XXXX | Zimbabwe | Middle Zambezi | 5 |
|  | *Oreochromis* cf. *mortimeri* | ZIM049 | PPZIM159.1 | XXXX | Zimbabwe | Middle Zambezi | 5 |
|  | *Oreochromis* cf. *mortimeri* | ZIM050 | PPZIM160.1 | XXXX | Zimbabwe | Middle Zambezi | 5 |
|  | *Oreochromis* cf. *mortimeri* | ZIM067 | PPZIM162.1 | XXXX | Zimbabwe | Middle Zambezi | 5 |
|  | *Oreochromis niloticus* (Linnaeus, 1758) | ZIM083 | PPZIM040.1 | XXXX | Zimbabwe | Middle Zambezi | 11 |
|  | *Oreochromis niloticus* (Linnaeus, 1758) | ZIM059 | PPZIM050.1 | XXXX | Zimbabwe | Middle Zambezi | 5 |
| *C. halli* (Price & Kirk, 1967) | *Coptodon rendalli* (Boulenger, 1897) | ZIM051 | PPZIM205.1 | XXXX | Zimbabwe | Middle Zambezi | 5 |
|  | *Oreochromis niloticus* (Linnaeus, 1758) | ZIM021 | PPZIM065_1 | XXXX | Zimbabwe | Middle Zambezi | 1 |
|  | *Oreochromis niloticus* (Linnaeus, 1758) | ZIM026 | PPZIM140_1 | XXXX | Zimbabwe | Middle Zambezi | 1 |
|  | *Oreochromis niloticus* (Linnaeus, 1758) |  |  | 203 (1) | DRC |  |  |
|  | *Oreochromis niloticus* x *mweruensis* |  |  | MG973075.1 (2) | DRC |  |  |
|  | *Oreochromis niloticus* (Linnaeus, 1758) |  |  | AJ920272.1 (3) | Ivory Coast |  |  |
|  | *Sarotherodon melanotheron* Rüppel, 1852 |  |  | AJ920271.1 (3) | Ivory Coast |  |  |
|  | *Sarotherodon galilaeus* (Linnaeus, 1758) |  |  | AJ920273.1 (3) | Ivory Coast |  |  |
|  | ? |  |  | KX869725.1 (4) | ? |  |  |
|  | *Oreochromis niloticus* (Linnaeus, 1758) |  |  | MH767392.1 (5) | Madagascar |  |  |
|  | *Sarotherodon galilaeus* (Linnaeus, 1758) |  |  | HE792790.1 (6) | Senegal |  |  |
|  | *Oreochromis niloticus* (Linnaeus, 1758) |  |  | MN709046.1 (7) | Egypt |  |  |
| *Cichlidogyrus acerbus* Dossou, 1982 | *Sarotherodon galilaeus*  (Linnaeus, 1758) |  |  | HE792780.2 (6) |  |  |  |
| *Cichlidogyrus aegypticus* Ergens, 1981 | *Coptodon guineensis*  (Günther, 1862) |  |  | HE792781.1 (6) |  |  |  |
| *Cichlidogyrus agnesi* Pariselle & Euzet, 1995 | *Coptodon guineensis*  (Günther, 1862) |  |  | AJ920286.1 (3) |  |  |  |
| *Cichlidogyrus arthracanthus* Paperna, 1960 | *Coptodon guineensis*  (Günther, 1862) |  |  | HE792783.1 (6) |  |  |  |
| *Cichlidogyrus attenboroughi* Kmentová, Gelnar, Koblmüller & Vanhove, 2016 | *Benthochromis tricoti* (Poll, 1948) |  |  | MH708153.1 (8) |  |  |  |
| *Cichlidogyrus berradae* Pariselle & Euzet, 2003 | *Coptodon rendalli* (Boulenger, 1897) |  |  | 182 (1) |  |  |  |
| *Cichlidogyrus bilongi* Pariselle & Euzet, 1995 | *Coptodon guineensis* (Günther, 1862) |  |  | AJ920287.1 (3) |  |  |  |
| *Cichlidogyrus brunnensis* Kmentová, Gelnar, Koblmüller & Vanhove, 2016 | *Trematocara unimaculatum* Boulenger, 1901 |  |  | MH708152.1 (8) |  |  |  |
| *Cichlidogyrus buescheri* Pariselle & Vanhove, 2015 | *Interochromis loocki* (Poll, 1949) |  |  | 363 (9) |  |  |  |
| *Cichlidogyrus casuarinus* Pariselle, Muterezi Bukinga & Vanhove, 2015 | *Bathybates minor* Boulenger, 1905 |  |  | KX007775.1 (10) |  |  |  |
| *Cichlidogyrus cirratus* Paperna, 1964 | *Oreochromis niloticus* (Linnaeus, 1758) |  |  | HE792784.1 (6) |  |  |  |
| *Cichlidogyrus consobrini* Jorissen, Pariselle & Vanhove in Jorissen, Pariselle, Huyse, Vreven, Snoeks, Volckaert, Chocha Manda, Kapepula Kasembele, Artois & Vanhove, 2017 | *Sargochromis mellandi*  (Boulenger, 1905) |  |  | 144 (1) |  |  |  |
| *Cichlidogyrus cubitus* Dossou, 1982 | *Coptodon guineensis* (Günther, 1862) |  |  | HE792785.1 (6) |  |  |  |
| *Cichlidogyrus digitatus* Dossou, 1982 | *Coptodon guineensis* (Günther, 1862) |  |  | HE792786.1 (6) |  |  |  |
| *Cichlidogyrus dossoui* Douëllou, 1993 | *Coptodon rendalli* (Boulenger, 1897) |  |  | 85 (1) |  |  |  |
| *Cichlidogyrus douellouae* Pariselle, Bilong Bilong & Euzet, 2003 | *Sarotherodon galilaeus*  (Linnaeus, 1758) |  |  | HE792787.1 (6) |  |  |  |
| *Cichlidogyrus dracolemma* Řehulková, Mendlová & Šimková, 2013 | *Hemichromis letournaeuxi* Sauvage, 1880 |  |  | HE792794.1 (6) |  |  |  |
| *Cichlidogyrus ergensi* Dossou, 1982 | *Coptodon guineensis* (Günther, 1862) |  |  | HE792788.1 (6) |  |  |  |
| *Cichlidogyrus falcifer* Dossou & Birgi, 1984 | *Hemichromis fasciatus*  Peters, 1857 |  |  | HE792789.1 (6) |  |  |  |
| *Cichlidogyrus flexicolpos* Pariselle & Euzet, 1995 | *Coptodon guineensis* (Günther, 1862) |  |  | AJ920283.1 (3) |  |  |  |
| *Cichlidogyrus gallus* Pariselle & Euzet, 1995 | *Coptodon guineensis* (Günther, 1862) |  |  | AJ920285.1 (3) |  |  |  |
| *Cichlidogyrus irenae* Gillardin, Vanhove, Pariselle, Huyse & Volckaert, 2012 | Gnathochromis pfefferi  (Boulenger, 1898) |  |  | KT692939.1 (10) |  |  |  |
| *Cichlidogyrus longicirrus* Paperna, 1965 | *Hemichromis fasciatus* Peters, 1857 |  |  | HE792791.1 (6) |  |  |  |
| *Cichlidogyrus mbirizei* Muterezi Bukinga, Vanhove, Van Steenberge & Pariselle, 2012 | *Oreochromis niloticus* x *mweruensis* |  |  | MG973076.1 (2) |  |  |  |
| *Cichlidogyrus nageus* Řehulková, Mendlová & Šimková, 2013 | *Sarotherodon galilaeus* (Linnaeus, 1758) |  |  | HE792795.1 (6) |  |  |  |
| *Cichlidogyrus njinei* Pariselle, Bilong Bilong & Euzet, 2003 | *Sarotherodon galilaeus*  (Linnaeus, 1758) |  |  | HE792792.1 (6) |  |  |  |
| *Cichlidogyrus nshomboi* Muterezi Bukinga, Vanhove, Van Steenberge & Pariselle, 2012 | *Boulengerochromis microlepis* (Boulenger, 1899) |  |  | 368 (9) |  |  |  |
| *Cichlidogyrus philander* Douëllou, 1993 | *Pseudocrenilabrus philander*  (Weber, 1897) |  |  | MG250200.1 (11) |  |  |  |
| *Cichlidogyrus pouyaudi* Pariselle & Euzet, 1994 | *Tylochromis intermedius* (Boulenger, 1916) |  |  | HE792793.1 (6) |  |  |  |
| *Cichlidogyrus schreyenbrichardorum* Pariselle & Vanhove, 2015 | *Interochromis loocki* (Poll, 1949) |  |  | 362 (9) |  |  |  |
| *Cichlidogyrus sclerosus* Paperna & Thurston, 1969 | *Oreochromis niloticus* (Linnaeus, 1758) |  |  | DQ537359.1 (12) |  |  |  |
| *Cichlidogyrus thurstonae* Ergens, 1981 | *Paretroplus lamenabe* Sparks 2008 |  |  | MH767395.1 (5) |  |  |  |
| *Cichlidogyrus tiberianus* Paperna, 1960 | *Coptodon guineensis* (Günther, 1862) |  |  | HE792796.1 (6) |  |  |  |
| *Cichlidogyrus tilapiae* Paperna, 1960 | *Hemichromis fasciatus*  Peters, 1857 |  |  | HE792797.1 (6) |  |  |  |
| *Cichlidogyrus vealli* Pariselle & Vanhove, 2015 | *Interochromis loocki* (Poll, 1949) |  |  | 356 (9) |  |  |  |
| *Cichlidogyrus yanni* Pariselle & Euzet, 1996 | *Coptodon guineensis* (Günther, 1862) |  |  | HE792798.1 (6) |  |  |  |
| *Cichlidogyrus zambezensis* Douëllou, 1993 | *Serranochromis macrocephalus* (Boulenger, 1899) |  |  | 375 (9) |  |  |  |
| *Scutogyrus bailloni* Pariselle & Euzet, 1995 | *Sarotherodon galilaeus*  (Linnaeus, 1758) |  |  | HE792799.1 (6) |  |  |  |
| *Scutogyrus longicornis* (Paperna & Thurston, 1969) | *Oreochromis niloticus* (Linnaeus, 1758) |  |  | HE792800.1 (6) |  |  |  |
| *Scutogyrus minus* (Dossou, 1982) | *Sarotherodon melanotheron* Rüppel, 1852 |  |  | HE792801.1 (6) |  |  |  |
| ***COI*** |  |  |  |  |  |  |  |
| *Cichlidogyrus chloeae* **sp. nov.** | *Oreochromis* cf. *mortimeri* | ZIM050 | PPZIM160.1 | XXXX | Zimbabwe | Middle Zambezi | 5 |
|  | *Oreochromis* cf. *mortimeri* | ZIM067 | PPZIM162.1 | XXXX | Zimbabwe | Middle Zambezi | 5 |
|  | *Oreochromis niloticus (*Linnaeus, 1758) | ZIM083 | PPZIM040.1 | XXXX | Zimbabwe | Middle Zambezi | 11 |
| *Cichlidogyrus halli* (Price & Kirk, 1967) | *Oreochromis niloticus (*Linnaeus, 1758) | ZIM029 | PPZIM019_1 | XXXX | Zimbabwe | Middle Zambezi | 2 |
|  | *Oreochromis niloticus (*Linnaeus, 1758) | ZIM021 | PPZIM065_1 | XXXX | Zimbabwe | Middle Zambezi | 1 |
|  | *Oreochromis niloticus (*Linnaeus, 1758) | ZIM026 | PPZIM140_1 | XXXX | Zimbabwe | Middle Zambezi | 1 |
| *Cichlidogyrus halli* (Price & Kirk, 1967) | *Oreochromis niloticus* x *mweruensis* |  |  | MG970255.1 (2) | DRC |  |  |
|  | *Oreochromis niloticus* (Linnaeus, 1758) |  |  | 203 (1) | DRC |  |  |
| *Cichlidogyrus sclerosus* Paperna & Thurston, 1969 | ? |  |  | JQ038226.1 (13) |  |  |  |
| *Cichlidogyrus zambezensis* Douëllou, 1993 | *Serranochromis jallae* (Boulenger 1896) |  |  | KT037411.1 (14) |  |  |  |
| *Cichlidogyrus philander* Douëllou, 1993 | *Pseudocrenilabrus philander*  (Weber, 1897) |  |  | MG288503.1 (15) |  |  |  |
| *Cichlidogyrus mbirizei* Muterezi Bukinga, Vanhove, Van Steenberge & Pariselle, 2012 | ? |  |  | MN905506.1 (16) |  |  |  |
| *Cichlidogyrus irenae* Gillardin, Vanhove, Pariselle, Huyse & Volckaert, 2012 | *Gnathochromis pfefferi*  (Boulenger, 1898) |  |  | KT037339.1 (14) |  |  |  |
| *Cichlidogyrus dossoui* Douëllou, 1993 | *Coptodon rendalli* (Boulenger, 1897) |  |  | 85 (1) |  |  |  |
| *Cichlidogyrus falcifer* Dossou & Birgi, 1984 | *Hemichromis stellifer* Loiselle, 1979 |  |  | 193 (1) |  |  |  |
| *Cichlidogyrus papernastrema* Price, Peebles & Bamford, 1969 | *Tilapia sparrmanii* Smith, 1840 |  |  | 80 (9) |  |  |  |
| *Cichlidogyrus quaestio* Douëllou, 1993 | *Coptodon rendalli* (Boulenger, 1897) |  |  | 83 (1) |  |  |  |
| *Cichlidogyrus thurstonae* Ergens, 1981 | *Oreochromis niloticus* (Linnaeus, 1758) |  |  | 214 (1) |  |  |  |
| *Cichlidogyrus tilapiae* Paperna, 1960 | *Oreochromis niloticus* (Linnaeus, 1758) |  |  | 208 (1) |  |  |  |
| *Scutogyrus gravivaginus* (Paperna & Thurston, 1969) | *Oreochromis mweruensis* Trewavas, 1983 |  |  | 65 (9) |  |  |  |

References: (1) (Jorissen et al., 2021), (2) (Vanhove et al., 2018), (3) (Pouyaud et al., 2006), (4) Francisco et al., Unpublished), (5) (Šimková et al., 2019), (6) (Mendlová et al., 2012), (7) Eldeep & Abdel Razik, Unpublished, (8) (Kmentová et al., 2018), (9) (Cruz-Laufer et al., 2021), (10) (Kmentová et al., 2016), (11) (Dos Santos, Unpublished results), (12) (Wu et al., 2007), (13) (Zhang et al., Unpublished), (14) (Vanhove et al., 2015), (15) (Igeh et al., 2017), (16) (Rong et al., Unpublished results)

### Fig. S1

Biplots of the PCAs plotting the second and third principal components PC2 and PC3: a PCA based on all measurements, b only measurements of the haptor, and c only measurements of the MCO. Each dot represents one specimen. Different colours represent different species i.e. *C. halli* and *C. chloeae* sp. nov. Ellipses are drawn at a confidence interval of 0.95. The contribution of the different measurements to the principal components are shown by arrows.

### Fig. S2

Maximum likelihood phylogenetic trees of specimens of *Cichlidogyrus* and *Scutogyrus* inferred from the 18S-ITS1 fragment (left) and *COI* fragment (right). Only well supported nodes (bootstrap values ≥ 0.85) are indicated by support values (in red). Scale bar indicates number of substitutions per site. Specimens of *C. chloeae* sp. nov. framed in green, specimens of *C. halli* framed in orange. Parasite labels and Genbank accession numbers of the included specimens can be found in Table S1.
